# Motor Cortex Modulates Ipsilateral Limb Movement Through a Direct Cortico-Cerebellar Circuit

**DOI:** 10.1101/2025.07.08.663625

**Authors:** Carmen Schäfer, Rui Oliveira Silva, Hana Hasanbegovic, Youri Adolfs, Jeroen Pasterkamp, Xiangning Li, Anan Li, Hui Gong, Chris De Zeeuw, Freek Hoebeek, Zhenyu Gao

## Abstract

Motor cortex is traditionally associated with control of contralateral limb movements via corticospinal and cortico-ponto-cerebellar pathways. However, the contribution of ipsilateral motor cortical outputs on motor control remains unclear. Here, we identify and characterize a distinct population of cortico-cerebellar (C-C) neurons in the motor cortex that form monosynaptic projections to the ipsilateral cerebellar nuclei. The C-C neurons receive preferential local motor cortical inputs and exhibit projection patterns distinct from cortico-pontine projecting neurons. Using in vivo imaging and optogenetic perturbations, we show that these neurons are active during locomotion and transitions of volitional movements. Disruption of the C-C projection severely affects the locomotion and balancing. Interestingly, the C-C pathway is selectively involved in the initiation and coordination of ipsilateral forelimb movements, without affecting contralateral movement kinematics. These findings shed light on a non-canonical cortico-cerebellar pathway that supports ipsilateral motor control, complementing the traditional control mechanisms of the cerebral cortex over the contralateral motor domains.

## Introduction

Goal directed voluntary movements are self-generated purposeful actions essential for daily behaviour. The brain integrates externally and internally generated information and generates coordinated motor outputs that control specific movements ^1–5^. This transformation engages distributed brain regions, which function as components of integrated, large-scale neural networks rather than isolated modules ^1,6–11^.

Among these regions, the motor cortex and cerebellum play foundational roles in generating and refining motor functions. The motor cortex is considered to be essential for preparing and generating motor commands ^12,13^. One of the prevailing views of motor control theory states that population activity of motor cortical neurons operates as a dynamical system ^14–18^, where population activity patterns drive movement execution. Although individual neurons may exhibit distinct activity patterns over time, movement-related states are encoded at the population level. These dynamics are essential for both movement preparation and execution ^18^.

The cerebellum, on the other hand, is widely recognized for its role in generating predictions and refining motor commands during motor control ^19–21^. By comparing intended motor commands with sensory feedback from executed movements, the cerebellum refines ongoing and future motor actions. This predictive function allows for fine-tuning adjustments that enhance movement accuracy and coordination ^19,20,22,23^. The communication between the motor cortex and cerebellum occurs primarily via the cortico-ponto-cerebellar pathway, wherein corticospinal neurons (CSNs) collateralize to pontine nuclei, relaying signals predominantly to the contralateral cerebellum ^24–32^. This anatomical organization positions CSNs as a key cortical output channel communicating with the cerebellum ^33–35^.

Voluntary muscle activation is primarily driven by lateralized motor commands originating from the contralateral primary motor cortex ^36,37^. Such lateralization of motor control is largely attributable to the distinct topography and projection patterns of the CSN descending pathways. The majority of motor cortical CSNs decussate at the level of the medullary pyramids and innervate premotor interneurons within the contralateral spinal cord, thereby facilitating the precise control of voluntary movements in the contralateral musculature ^36,38–44^. The cerebellar outputs, i.e. the cerebellar nuclei (CN) neurons on the other hand, descend via contralateral red nucleus to influence ipsilateral spinal circuits ^45,46^, and predominantly affect ipsilateral movements ^47^. Therefore, it is believed that movements are coordinated by integrative activities of the contralateral motor cortex and the ipsilateral cerebellum, coordinated by the contralateral dominant cortico-cerebellar communication loops ^28,30,44,45^.

Yet, emerging evidence demonstrates that motor cortical activity is not exclusively dedicated to contralateral motor output. Ipsilateral motor-related activity in motor cortex has been documented in both human and animal studies ^48–55^. A substantial proportion of motor cortical neurons are engaged during ipsilateral limb movements ^48,54^. Interestingly, these neurons exhibit distinct encoding of ipsilateral and contralateral movements, often occupying orthogonal representational subspaces ^53^. Such neuronal representation may ensure that ipsilateral motor representations do not interfere with contralateral motor commands ^54,56^, but the functional importance of the motor cortical outputs for the ipsilateral side of the body remain poorly understood.

In this study, we identify an unconventional direct cerebro-cerebellar pathway that bypasses the traditional cortico-ponto-cerebellar circuit, enabling monosynaptic communication from the motor cortex to the ipsilateral cerebellum. We show that activity in this pathway controls both ongoing locomotion and skilled reaching movements. Disrupting this direct ipsilateral cerebro-cerebellar communication through photo-perturbation halts locomotion and disturbs the timing of ipsilateral but not contralateral forelimb movements, highlighting its unique role in facilitating ipsilateral motor coordination. Together, these findings reveal a novel cortico-cerebellar circuit with distinct laterality and functional significance.

## Results

### Anatomical characterisation of direct cortico-cerebellar pathway

The CN neurons receive afferent inputs from the cerebral cortex primarily through mossy and climbing fiber pathways, which originate from various relay nuclei within the brainstem (Fig. 1a) ^30,57^. We first sought to identify whether the CN receive inputs from cerebrum via other cerebro-cerebellar pathways. We performed mono-synaptic retrograde tracing by injecting retrograde adeno-associated virus (AAVretro) encoding *Cre*-recombinase in the CN of the *Ai14* reporter mice (Fig. 1b) ^58^. Bilateral injections covering the entire CN regions revealed long-distance CN-projecting neurons in various brain regions. Surprising we observed very abundant neurons that originate from the cerebral cortex (Fig. 1c-d, Extended Data Fig. 1a-c). These neurons were termed as the direct cerebro-cerebellar projecting (C-C) neurons.

**Figure 1.**
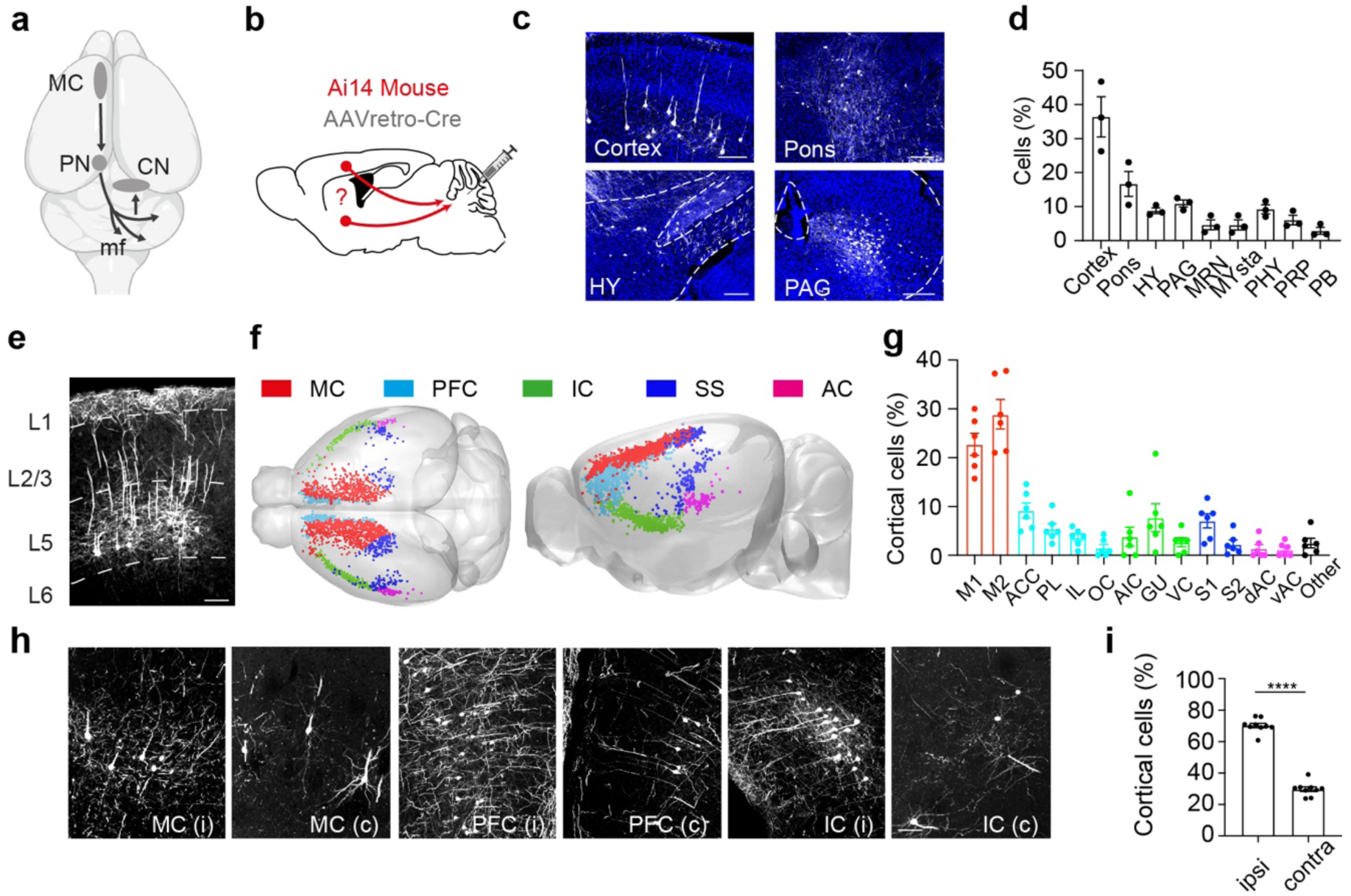
Anatomical characterization of the cortico-cerebellar pathway. a. Schematic illustration of the laterality of the projections between MC and cerebellum. MC predominantly controls the contralateral cerebellum via the mossy fiber (mf) projections that are relayed via the ipsilateral pontine nuclei (PN). The CN integrate outputs from the cerebellar cortex. b. Experimental procedure for retrograde tracing from CN. The conditional expression of tdTomato identified cell populations in extra-cerebellar input areas that complement projections from pontine nuclei (PN), inferior olive (IO) and vestibular nuclei. c. Images illustrating extra-cerebellar brain areas that innervate cerebellar nuclei. The most prominent input areas are cerebral cortex, pons, hypothalamus (HY) and periaqueductal gray (PAG). Scale bars are 200 µm. d. Quantification of the regions and neurons that project to CN. Hypothalamus (HY), periaqueductal gray (PAG), midbrain reticular nucleus (MRN), medulla (MY), perihypoglosssal nuclei (PHY), nucleus prepositus (PRP) and parabrachial nucleus (PR) e. Image of CN-projecting neurons in the layer 5 of MC. Scale bar is 100 µm. f. Distribution of cortico-cerebellar projecting neurons in the cerebral cortex of an iDISCO cleared brain. Motor Cortex (MC), prefrontal cortex (PFC), insular cortex (IC), sensory cortex (SS) and auditory cortex (AC). g. Distribution of cortico-cerebellar projecting neurons. Primary motor cortex (M1), secondary motor cortex (M2), anterior cingulate cortex (ACC) prelimbic cortex (PL), infralimbic cortex (IL), orbitofrontal cortex (OC), agranular insular cortex (AIC) gustatory cortex (GU), visceral cortex (VC), primary sensory cortex (S1), secondary sensory cortex (S2), dorsal auditory cortex (dAC), ventral auditory cortex (vAC). N = 6. h. Images of layer 5 pyramidal neurons in ipsi (i) and contralateral (c) motor cortex (MC), prefrontal cortex (PFC) and insular cortex (IC). Scale bar 50 µm. i. Ipsilateral and contralateral distribution of cortico-cerebellar projecting neurons. N = 9, paired t-test, p < 0.0001. All values represent the mean ± sem.

We performed iDISCO based brain clearing and light-sheet whole brain imaging to visualize the distribution of CN-projecting neurons across the cerebral cortex ^59,60^ (Fig. 1e-g, Supplementary Video 1). Cell registration showed that the C-C neurons were mostly localized to the motor cortical regions (MC) and to a lesser extent in the prefrontal, insular, and sensory cortices (Fig. 1e-g). Hardly any C-C neurons were observed in other cerebral cortical regions. This specific cortical distribution of C-C neurons stands in marked contrast to that of the PN-projecting corticospinal neurons (CSN), which are widely distributed throughout cerebral cortex ^61,62^. Unilateral viral tracing from the CN revealed that the majority of C-C input neurons (70.3 +/- 1.5%) localized to the cerebral cortex ipsilateral to the cerebellar injection site (Fig. 1h-i, Extended Data Fig. 1 a-c).

### Whole-brain mapping of cortico-cerebellar neurons reveals unique projection patterns

As the majority of C-C neurons originate from motor cortical regions, we examined the downstream projections of the motor cortical C-C neurons. We combined the cell-type specific viral labeling strategy with whole-brain visualization of quantification using fluorescence micro-optical sectioning tomography (fMOST) ^61,63,64^ to map the axonal projections throughout the entire brain. We injected AAVretro encoding *Cre*-recombinase to CN and cre-dependent GFP-expressing virus in the MC to selectively label the C-C neurons in the MC (Fig. 2a, Extended Data Fig. 2a). We reconstructed the complete axonal projection patterns of 29 C-C neurons from 3 mice and quantified their whole brain projectome in detail (Fig. 2b-f, Extended Data Fig. 2b,c). All C-C neurons showed elaborate axonal collateralization to cortical and subcortical regions (Fig. 2d-f). The most prominent downstream targets included the cerebral cortex, striatum, and midbrain areas such as zona incerta, superior colliculus, midbrain reticular nucleus, periaqueductal gray, pontine reticular nucleus, pontine nuclei, and inferior olive (Fig. 2d-f, Extended Data Fig. 2). Single axon quantification confirmed a strong ipsilateral preference of the C-C axonal projections, apart from the innervation to medullary nuclei, which was in many cases equal for the ipsilateral and contralateral sides (Fig. 2f). Unsupervised clustering analysis of reconstructed neurons revealed that all these neurons fell within the same cluster, implying a highly unique but consistent projection feature of the C-C neurons (Fig. 2f). The projections of C-C neurons to the spinal cord crossed to the contralateral side via the pyramidal tract (Fig. 2g). These data indicate that the C-C neurons are a specific sub-group of CSNs.

**Figure 2.**
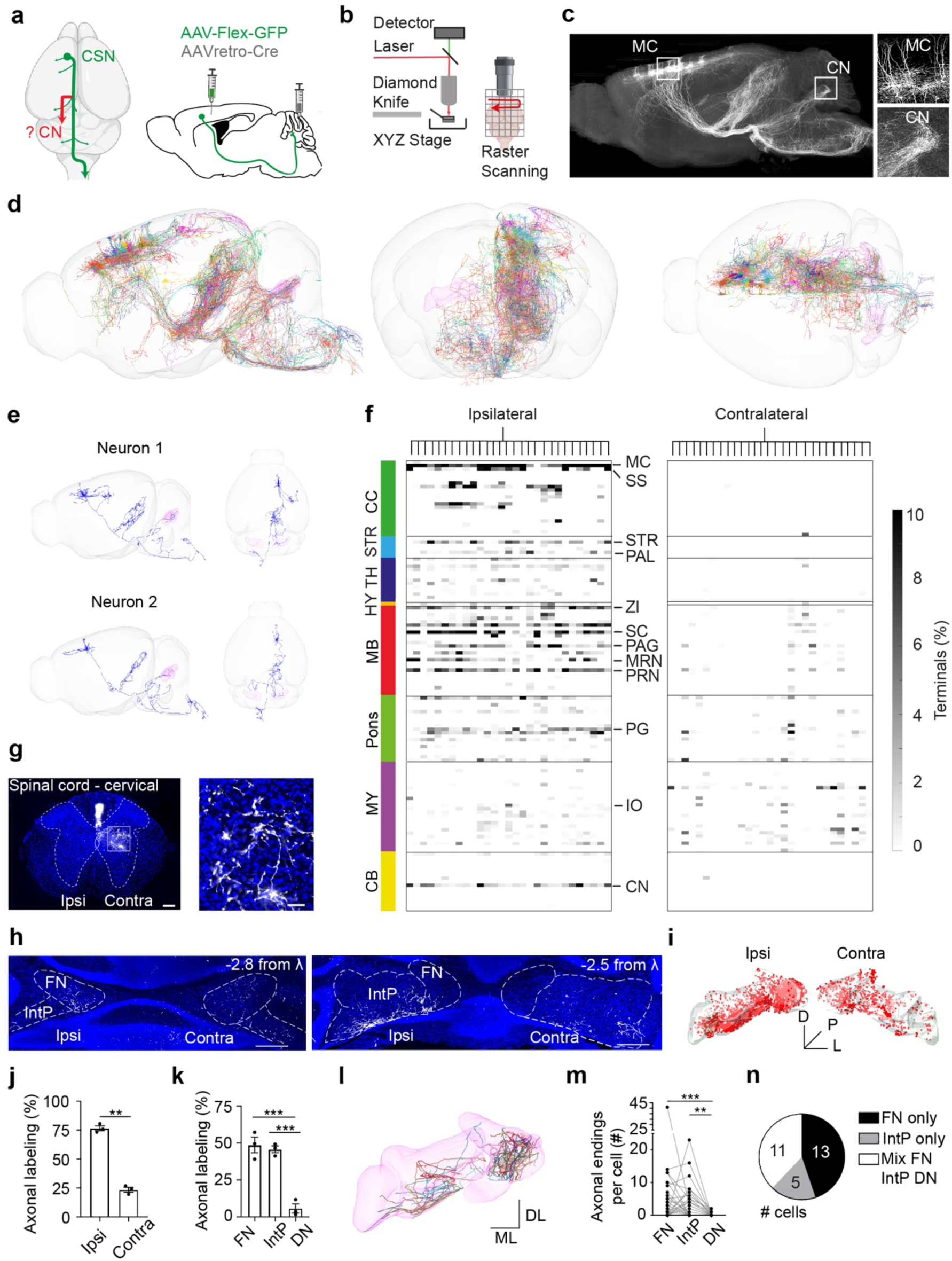
Brain-wide projection patterns of the C-C neurons. a. Targeted retrograde tracing strategy to label C-C neuron projections. b. Schematic depiction of the fMOST imaging technique. c. Maximum projection of fMOST images showing the whole brain projection patterns of C-C neurons, highlighting the direct axonal projections to CN. Higher magnification views of C-C neuron somata in the motor cortex and axon terminals in CN. d. Sagittal, coronal and horizontal views of all 29 C-C neurons and their projection patterns throughout the whole brain. Cells are registered to CCFv3, rendered in 3D, overlaid and randomly colored. N = 3. e. Sagittal and horizontal view of the complete axonal projection patterns of 3 example C-C neurons. f. Heatmap showing the fraction of total terminal endpoints of the C-C neuron projection trajectory in all relevant downstream targets of the 29 C-C neurons. Columns represent individual cell; rows represent target areas ordered by the major brain divisions from the Allen CCF. The dendrogram on top shows the hierarchical cluster, indicating high similarity across all reconstructed cells. g. Images of C-C neuron innervation in spinal cord. h. Images of motor cortical innervation in FN and IntP. Scale bar: 500 µm. i. Distribution of C-C terminals in CN. j. The motor cortical innervation in the ipsilateral and contralateral CN. N = 3, paired t-test, p = 0.0062. k. Distribution of C-C terminals in FN, IntP and DN. N = 3, Ordinary one-way ANOVA with Bonferroni correction, FN vs. DN: p = 0.0006, IntP vs. DN: p = 0.0009. l. The terminal endings of the 29 reconstructed C-C neurons. m. Quantification of number of terminal endings in FN, IntP and DN. N = 3, Friedman test with Dunn’s test for multiple comparisons, FN vs. DN: p = 0.0001, IntP vs. DN: p = 0.0051. n. The proportions of C-C neurons that innervate FN, IntP or a mix of all three CN nuclei. All values represent the mean ± sem.

We next characterized the axonal innervation patterns in the CN. The C-C axon terminals were registered to the Allen CCF to quantify their axonal projection patterns in the CN (Fig. 2h-n). The innervation was most dominant in the ipsilateral Fastigial (FN) and Interposed Nucleus (IntP), while it remained largely absent in the Dentate Nucleus (DN, Fig. 2k-n, Extended Data Fig. 3a,b). Quantification of the axon innervation patterns from individual C-C neurons revealed that 45% (13/29) neurons innervated exclusively FN, 38% (11/29) innervated both FN and IntP, and 17% (5/29) innervated exclusively IntP (Fig. 2l-n, Extended Data Fig. 3a,b). Injections of AAVretro virus expressing RFP, GFP, and Cholera Toxin Subunit B (CTB) into different CNs (Extended Data Fig. 3c-d) confirms the proportion of C-C neurons (Extended Data Fig. 3e-g). These results collectively showed that C-C neurons selectively target specific sub-regions of the cerebellar nuclei.

### Complementary organizations of the monosynaptic C-C and the disynaptic cortico-ponto-cerebellar pathways

The most prominent pathway mediating communication between MC and cerebellum is the cortico-ponto-cerebellar pathway ^26,30,31,65^. More specifically the CSNs of the MC collateralize to the basal pontine nuclei, which in turn project to the cerebellum via mossy fibers. We next sought to characterize the common and distinct projection features of the monosynaptic C-C and disynaptic cortico-ponto-cerebellar pathways^30,31,65^.

We applied a dual retrograde labeling approach in which we delivered AAVretro virus expressing GFP to PN to label CSNs and AAVretro virus expressing RFP to CN to label C-C neurons (Extended Data Fig. 4a). Double labelled neurons confirmed that C-C neurons are a specific subpopulation of CSNs (Extended Data Fig. 4a-c). To further investigate the topography of cortico-ponto-cerebellar projections, we injected an anterograde trans-synaptic AAV1 virus encoding *Cre*-recombinase ^31,65^ in the MC and AAV9 expressing *Cre*-dependent GFP in the pontine nuclei. The CSN-targeted pontine neurons were selectively labelled with this approach, and their MF terminals were observed both in cerebellar cortex and CN (Fig. 3a-c). MC-targeted PN-MF boutons were primarily found in the hemispheres and posterior vermal regions of the cerebellar cortex (Fig. 3d, Extended Data Fig. 4d-e). We found that unilateral CSN neurons target both the ipsi- and contralateral hemispheres of the cerebellar cortex, but with a contralateral preference (Fig. 3d-f). Interestingly, co-registering the MC-MF and C-C axonal collaterals revealed largely non-overlapping termination patterns in the CN (Fig. 3d-f). The C-C pathways targeted mainly the FN and IntP ipsilateral to MC, whereas the MC-MF collaterals in CN occupied the contralateral IntP and DN (Fig. 3g). These results illustrate that the monosynaptic C-C pathway connects to the cerebellar regions that are largely complementary to those of the disynaptic MC-MF pathways.

**Figure 3.**
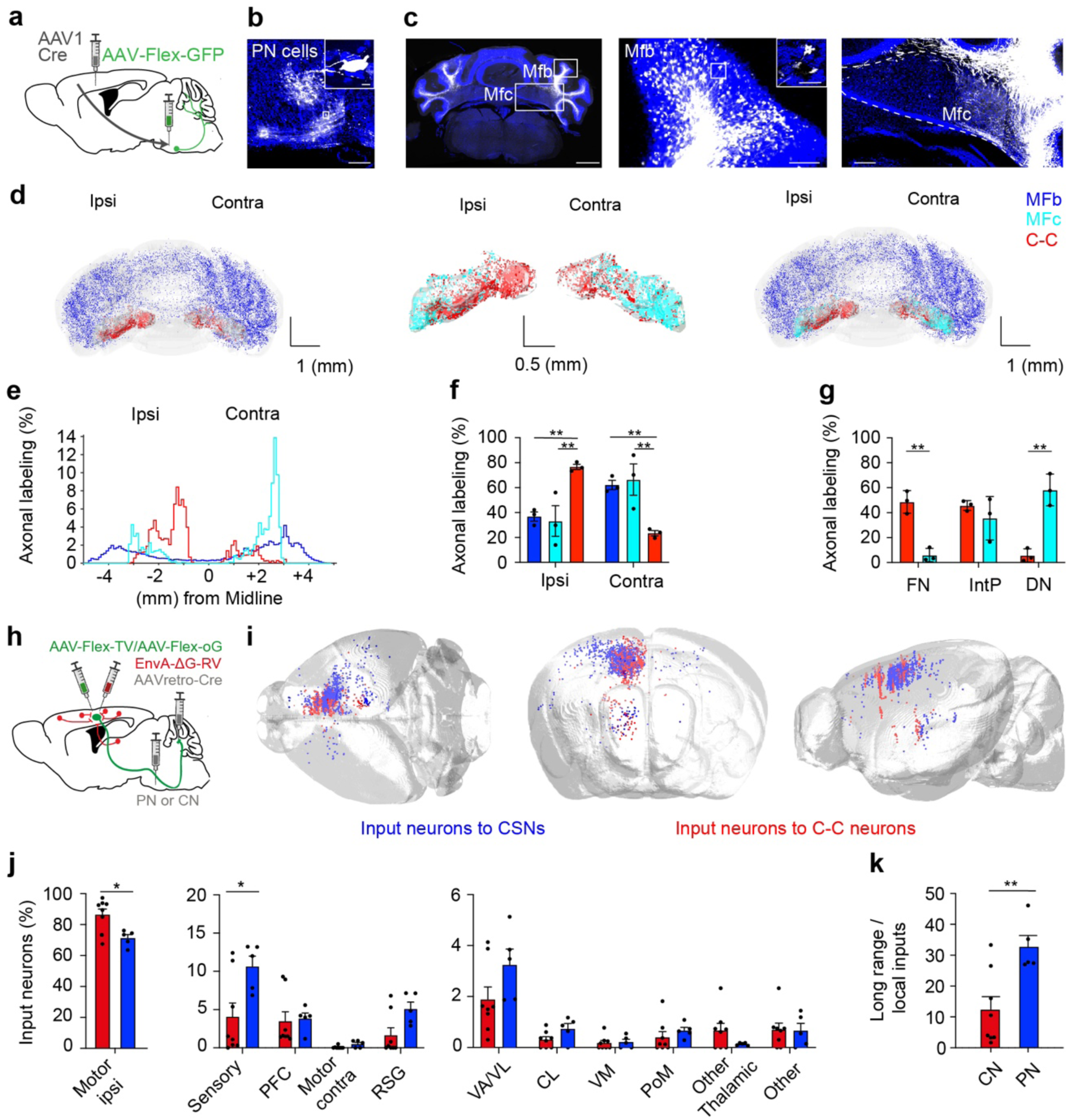
Complementary organizations of the C-C and cortico-ponto-cerebellar pathways. a. Transsynaptic viral labeling strategy for labelling the cortico-pontine MF pathway. b. Example PN neurons that receive MC input. Scale bar 100 µm. Scale bar zoom in: 20 µm. c. Left: cerebellar section showing mossy fiber boutons (Mfb) in the cerebellar cortex and mossy fiber collaterals (Mfc) in CN, Scale bar: 500 µm. Middle: zoom in image showing Mfb in the cerebellar cortex (scale bars: 200 and 20 µm). Right: zoom in image showing the mossy fiber collaterals (Mfc) in CN, scale bar: 200 µm. d. The Mfb in the cerebellar cortex, Mfc in the CN and the C-C neuron projections to CN are reconstructed and rendered in the 3D volume of the cerebellum. N = 3. e. Distribution of Mfb, Mfc and MCc axonal labeling along the medio-lateral axis of the cerebellum. N = 3. f. The relative innervation of Mfb, Mfc and MCc in the ipsilateral and contralateral side of the cerebellum. N = 3, two-way ANOVA with Tukey’s multiple comparison test, Mfb vs. MCc ipsi: p = 0.0077, Mfc vs. MCc ipsi: p = 0.0043, Mfb vs. MCc contra: p = 0.0093, Mfc vs. MCc contra: p = 0.0045. g. Quantification of Mfc and MCc innervation in FN, IntP and DN. Multiple t-test with Holm Sidak correction, FN: p = 0.007, DN: p = 0.007. h. Scheme of monosynaptic rabies viral tracing strategy for mapping input neurons to C-C neurons and CSNs. i. The distribution of input neurons to C-C neurons (red) and CSNs (blue) in two example brains overlaying on the Allen CCF. j. Distribution of inputs neurons to C-C neurons (N = 8) and CSNs (N = 5) from local cortical, long-range cortical and subcortical brain areas. Multiple t-tests with Holm-Sidak correction: Number of t-tests: 11, Motor ipsi: p < 0.000001, Sensory: p = 0.00668. k. The ratio between local and long-range cortical inputs for C-C neurons and CSNs. Unpaired t-test, p = 0.0069.

Given the distinct projection features of the C-C neurons, we next investigated which brain regions provide inputs to these C-C neurons. We performed projection-based monosynaptic retrograde rabies tracing to map the local and distant presynaptic neurons that project to either the C-C or the typical CSN neurons (Fig. 3h) ^66^. AAVretro-*Cre* virus was injected in either the C-C recipient CN region or the CSN recipient pontine region, while *Cre*-dependent TVA and rabies glycoprotein (oG) AAVs were injected in the MC. Subsequent infection with G deleted rabies virus selectively labeled C-C and CSN starter neurons, as well as their mono-synaptically connected presynaptic input neurons (Extended Data Fig. 4f-h). The number of starter cells and input cells was graded proportionally (Extended Data Fig. 4i). Retrograde rabies tracing revealed presynaptic neurons from various brain areas including local motor cortical networks, long-range cortical areas and long-range subcortical areas in the thalamus (Fig. 3i,j, Extended Data Fig. 4j). C-C neurons received almost exclusively inputs from the local motor regions and a lower proportion from other cortical regions, compared with the distribution of input neurons presynaptic to CSN neurons (Fig. 3k). These anatomical data suggest that the C-C pathway may serve as a local hub to integrate motor cortical signals and funnel them directly to the cerebellar output stations, implying their potential involvements in modulating movements.

### Involvements of the C-C Pathway during naturalistic volitional movements

Given that activation of MC CSN is considered the main motor command to drive action ^13,38,41,67,68^, we asked whether the C-C pathway may be involved in the control of volitional movements. We injected AAVretro-*Cre* virus in the CN and *Cre* dependent GCamP7f virus in the MC to monitor activity of C-C neurons. We tracked the activity of these neurons using one-photon Ca^2+^ imaging in head restrained mice running on a horizontal treadmill (Fig. 4a-c). The imaging window was centred on the caudal forelimb areas (CFA, Fig. 4d). Interestingly, we observed that activities of distinct C-C neurons were tuned for specific behavioural states. The first group of C-C neurons was preferentially activated during the active running phase (Fig. 4e), whereas another group of C-C neurons was active during the resting phase and mostly silent during running (Fig. 4f). Among all the recorded neurons, 11.7% of the C-C neurons preferentially active during resting epoch (resting-tuned neuron), while 25.3% of the neurons had preferential modulation during locomotion (locomotion-tuned C-C neurons, Fig. 4g,h). The activation of these neurons typically occurred during the locomotion-to-rest or rest-to-locomotion transitions (Fig. 4i,j). These data suggest that C-C neurons are likely to convey action-related representations, as well as the transition of locomotion states.

**Figure 4.**
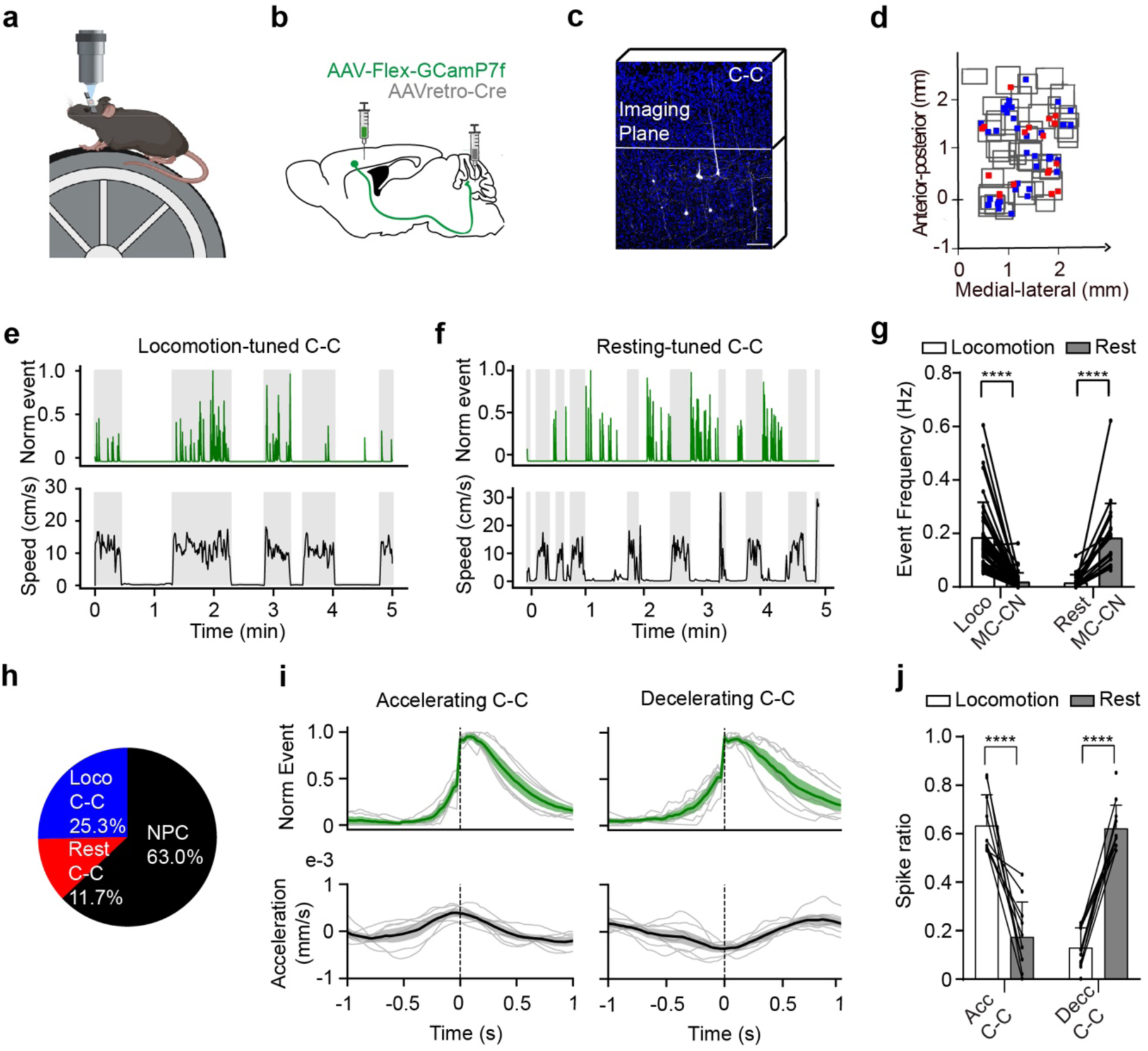
Involvements of the C-C Pathway in locomotion. a. Schematic diagram of the treadmill behavioural setup. b. Viral labelling strategy. c. Example C-C neurons and imaging plane. d. The imaging regions (gray squares) and cell locations (red and blue dots). Red dots represent resting-tuned C-C neurons and blue dots represent locomotion-tuned C-C neurons. e. Example calcium imaging events and locomotion speed for a locomotion tuned C-C neuron. f. Example calcium imaging events and locomotion speed for a resting tuned C-C neuron. g. Event frequency of locomotion- and resting-tuned C-C neurons during locomoting and resting epochs. Resting was defined as periods longer than 3 seconds without locomotion. N = 6, Wilcoxon test, **** p < 0.0001. h. Pie chart showing the proportions of locomotion-, rest- and non-modulating (NM) C-C neurons. i. Calcium events of C-C neurons precede both the locomotion-to-rest (deceleration) and the transition from rest-to-locomotion (acceleration) transitions. j. Quantification of the spike ratio in cells that modulate with deceleration and acceleration. Wilcoxon test, **** p < 0.0001.

### C-C pathway is crucial for coordinating volitional movements

We hypothesised that the C-C neurons activity during locomotion might be important for locomotion to coordinate the continuous adjustments of posture, body positioning and limb movements ^47,69–71^. To illustrate the functional relevance of the C-C pathway for locomotion, we expressed eOPN3, an inhibitory rhodopsin suitable for persistent terminal inhibition ^72^, in the C-C neurons. Optic fibers were implanted in the CN on both sides of the cerebellum (Fig. 5a). We selectively inhibited the C-C collaterals inside the CN and examined the impacts of terminal inhibition on voluntary locomotion (Fig. 5a,b, Extended Data Fig. 5a-c). We used high-speed video recording and Deeplabcut based tracking of limb positions and the nose to monitor the displacements of different body parts while the mouse was running through a glass hallway ^73^ (Fig. 5b). Bilateral terminal inhibition in running mice effectively slowed down the running speed of the mice, as indicated by the swing velocity of the forepaw, until they reached full stop (N = 8, Fig. 5c-g, Supplementary Video 2). When we analysed the basic stride parameters during control and after optogenetic inhibition, we found that stance time significantly increased and the swing length decreased (Fig. 5h-k), which is in line with the decreased running velocity. Rather strikingly, after reaching the full stop, a great effort was required from the mouse to re-initiate movements (Supplementary Video 2). As during each stride cycle the diagonal pair of fore and hindlimbs moves together and alternates with the other pair ^73,74^, we next tested whether optogenetic inhibition of unilateral C-C projection could bias the stride symmetry. We postulated that since C-C neurons target preferentially ipsilateral CN, the ipsilateral limb might be preferentially affected. Indeed, we found that the hindpaw ipsilateral to the photo-inhibition side had consistent delays in phase in relation to the reference forepaw, whereas the timing of the contralateral paw was unaffected (Fig. 5l-o, Extended Data Fig. 6a-l). To evaluate how this shift in phase synchrony affects posture and body positioning, we challenged the mice to walk over an elevated balance beam (Fig. 5p). Photo-inhibiting the C-C terminals in the CN induced hindpaw slip while walking on the beam (Fig. 5q), we analysed the laterality of hindpaw slip in control trials as well as in trials with left or right photo-inhibition. In control trials mouse slips were equally distributed between the left and right hindpaw (Fig. 5r). Consistent to the effects of ipsilateral inhibition during locomotion, Inhibiting C-C terminals in the left CN resulted in delays in stride phase of left limbs, pushing mice to fall to the right side. Inhibiting the right side of C-C terminals affected phase of right limbs and made mice to fall to the left (Fig. 5s,t, Supplementary Video 3). These results revealed a crucial role of C-C pathways in coordinating stride cycle and maintaining balance when mice were challenged to walk on the balance beam.

**Figure 5.**
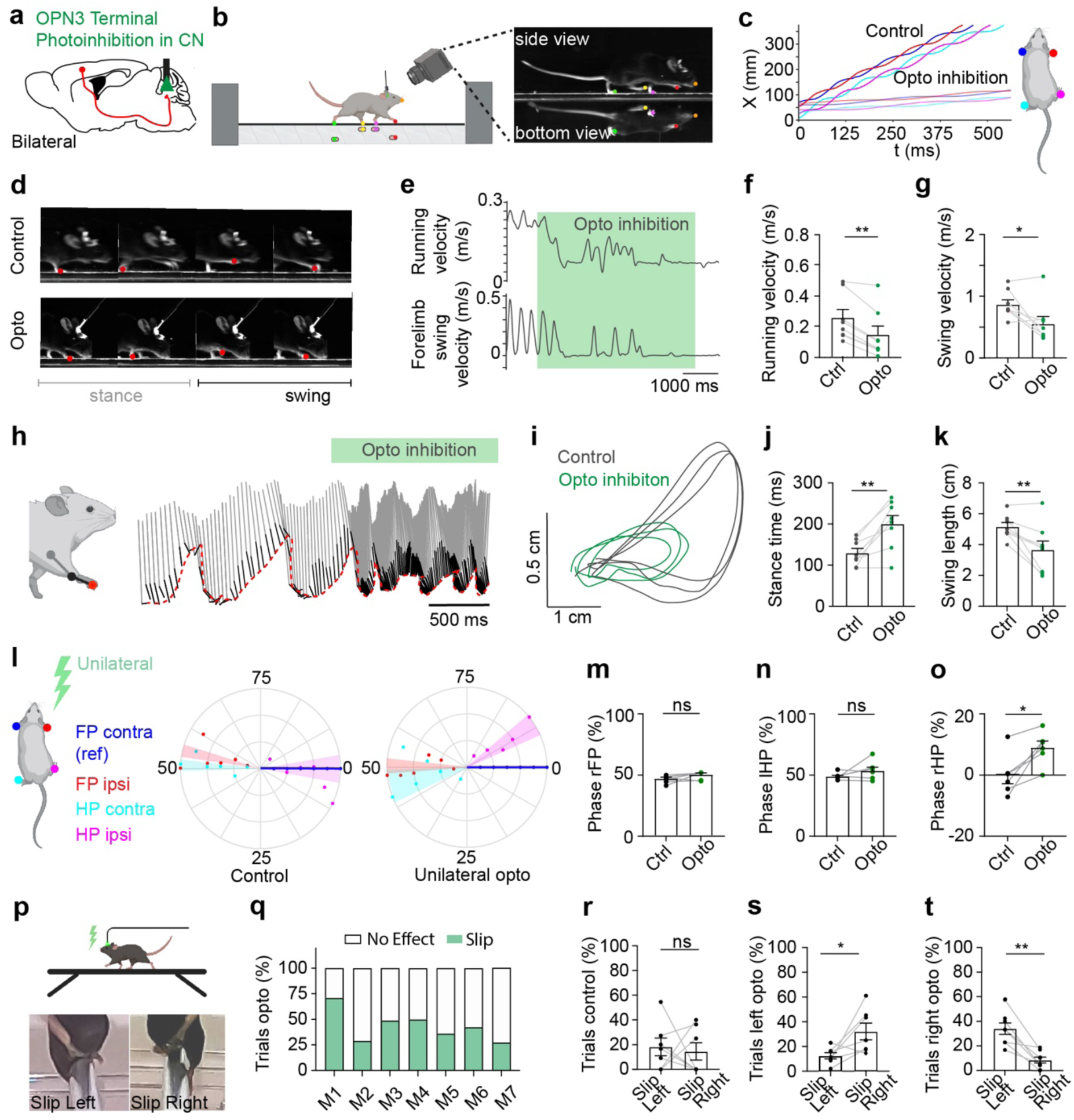
C-C pathway is crucial for coordinating volitional movements. a. Strategy for OPN3 photo-inhibition of C-C terminals in CN. b. Illustration showing a mouse running in Locomouse setup and DeepLabCut motion tracking. c. Continuous forward trajectories (x-position vs. time) for fore- and hindlimbs during control and photo-inhibition (opto) trials. Mouse inset illustrates colour codes for paws. d. Time series of stance and swing phases during control and photo-inhibition trials. The red dot depicts the position of the right forepaw. e. Running and swing velocity of the left forepaw (lFP) before and after photo-inhibition. f. Decreased running velocity after photo-inhibition. N = 8, Wilcoxon test, p = 0.0078. g. Decreased swing velocity after photo-inhibition. N = 8, Wilcoxon test, p = 0.0391. h. Color-coded stick diagram which depicts shoulder (gray), ancle (black) as well as paw endpoint (red) and represents forelimb movement before and after photo-inhibition. The red dash line depicts the forward trajectory of the forepaw. i. Trajectory of the forelimb movement during control (gray) and photo-inhibition (green). j. Quantification of the stance time during control and photo-inhibition. N = 8, Paired t-test, p = 0.0038. k. Quantification of the swing length during control and opto-inhibition. N = 8, Paired t-test, p = 0.0089. l. Polar plots indicating the phase of the step cycle in which each limb enters stance, aligned to stance onset of forepaw contralateral to stimulation (here left forepaw (lFP)) for control and unilateral photo-inhibition. Each dot represents one mouse and the shaded area represents sem. N = 6. Right forepaw (rFP), left hind paw (lHP) and right hind paw (rHP). m. Phase timing for the rFP stance onset. N = 6, Wilcoxon test, p = 0.1562. n. Same as m but for the lHP. N = 6, Wilcoxon test, p = 0.4375. o. Same as m but for the rHP. N = 6, Wilcoxon test, p = 0.0312. p. Balance beam experiment and photographs depicting slips of the left and right hindpaws. q. Percentage of trials with hind paw slips following unilateral photo-inhibition. M1-7 indicate individual mice. r. Percentage of slips in control trials. N = 7, paired t-test, p = 0.7636 s. Same as r but for the trials with C-C terminal inhibition in the left CN. N = 7, paired t-test, p = 0.0397. t. Same as r but for trials with C-C terminal inhibition in the right CN. N = 7, paired t-test, p = 0.0023.

To examine whether direct activation C-C pathway affects cerebellar output activity, we used multichannel silicone probes to record single-unit activity from CN neurons while photo-activating the channelrhodopsin-2 (ChR2) expressing C-C terminals in awake mice (Extended Data Fig. 7a-b). Activating C-C terminals facilitated CN activity dependent on the frequency of stimulation (Extended Data Fig. 7c-e), confirming that C-C pathway transmit an excitatory signal to CN. To investigate whether activation of C-C axons in the CN had effect on the postural control, we activated the C-C terminals while mice were moving on the balance beam (Extended Data Fig. 8a-c). Photo-activation of C-C terminals resulted in a significantly higher number of slips (Extended Data Fig. 8d-e). Consistent with the photo-inhibition experiments, unilateral activation of C-C terminals affected the limb movements of ipsilateral side, making mice to slip to the contralateral side (Extended Data Fig. 8f-h, Supplementary Video 3). Interestingly, photo-activating C-C terminals in awake but resting mice did not induce noticeable limb movements (Extended Data Fig. 7f-i), suggesting that the C-C pathway coordinates movements but does not drive them. Taken together these data indicate that C-C pathway conveys movement state-dependent information that is essential for organising ipsilateral movements.

### C-C Pathway contributes to the coordination of volitional forelimb reaching, but not to the control of reaching kinematics

It is well known that CSNs are predominantly involved in the initiation and execution of reaching of the contralateral forelimb ^12,32,75–79^, whereas selective disruption of the disynaptic cortico-ponto-cerebellar communication effects fine-tuning of reach kinematics ^80^. Our photoinhibition experiments during locomotion and balance beam identified ipsilateral effects of the C-C pathway, we next took the advantage of unimanual forelimb reaching movements to further clarify the lateralized effects of the C-C pathway on forelimb movements (Fig. 6a). During the task, a tone that acted as a go-cue was followed by a 2-second delay period and subsequently by the appearance of a water droplet ^75^. Mice were trained to perform volitional forelimb reaching to retrieve water within the 8-second reach epoch (Fig. 6a).

**Figure 6.**
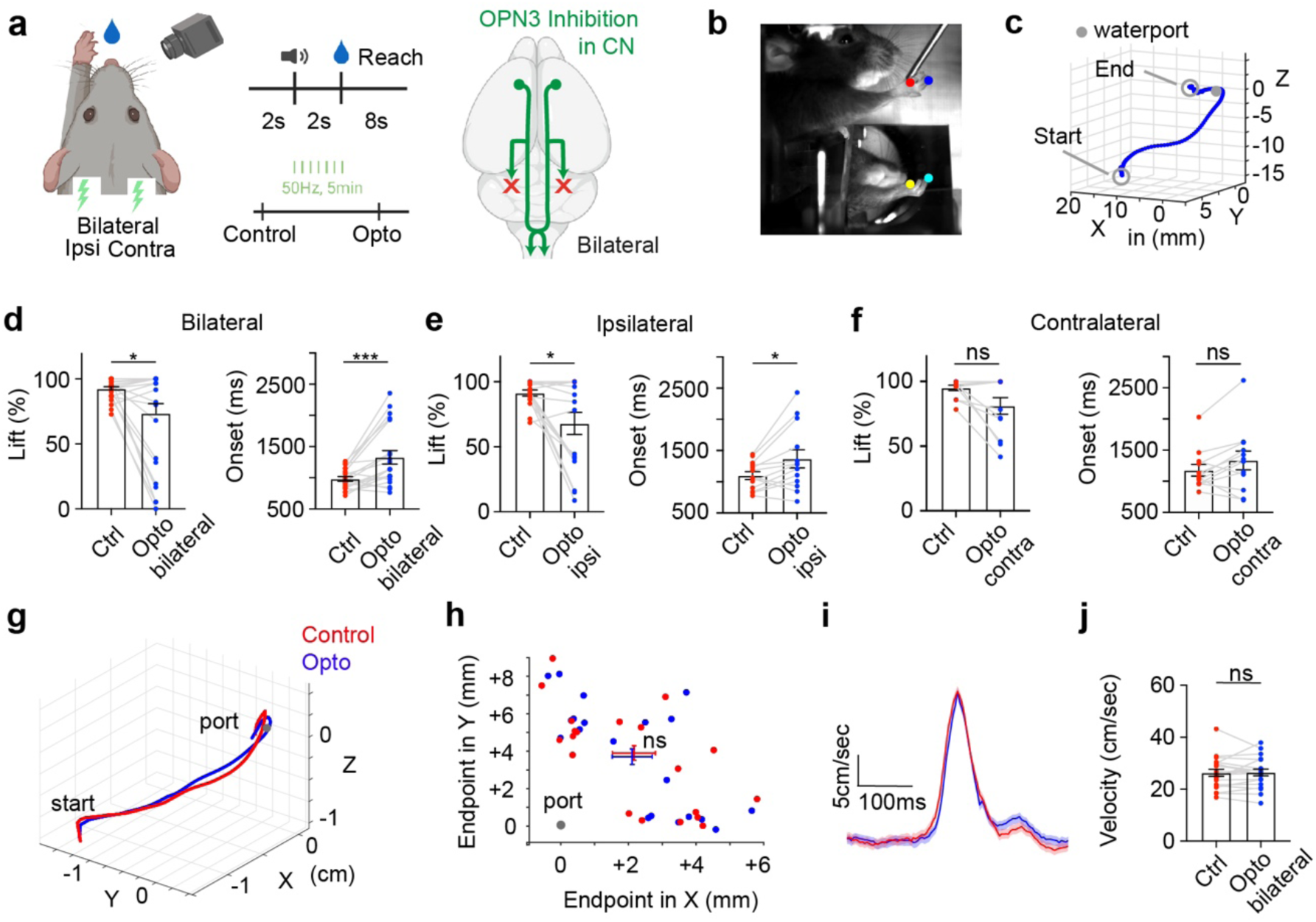
Photo-inhibition of C-C pathway affects the initiation but not the kinematics of forelimb reaching. a. Illustration of water droplet reaching paradigm and photo-inhibition strategy. b. Motion tracking of the forelimb coordinates with side and bottom views. c. 3D reconstruction of the paw reaching trajectory. d. Left: decreased lift probability after bilateral photo-inhibition of C-C terminals in the CN. N = 7, n = 25, p =0.0432. Right: delayed reach onset timing after bilateral inhibition of C-C terminals in the CN. N = 7, n = 20, p = 0.0003. e. Same as d, but for ipsilateral photo-inhibition. Decreased lift probability n = 17, p = 0.0134. Delayed reach onset timing. n = 13, p = 0.0181. f. Same as d, but for contralateral photo-inhibition. n = 12. P >0.05. g. Example reaching trajectories in control (red) and photo-inhibition (blue) trials. h. Distribution of the reaching endpoints in control and bilateral photo-inhibition trials. N = 7, n = 20, paired t-test, in X: p = 0.6043, in Y: p = 0.3299. i. Reaching velocy remains unaffected after bilateral silencing. Average across all sessions. Shading depicts sem. j. Quantification of reaching velocity for all sessions. N = 7, n = 20, paired t-test, p = 0.9010. All values represent the mean ± sem.

Once the mice became experts in the volitional reaching task, we tracked detailed movements of the forelimb (Fig 6b,c). The forelimb movements were video recorded and x-, y-, and z-coordinates of forepaw were extracted using the DeepLabCut behavioural analysis package ^81,82^. We first expressed eOPN3 in the C-C neurons and inhibited C-C terminals bilaterally (Fig. 6a). Bilateral optogenetic inhibition of the C-C terminals decreased the lift probability to less than 50% in 3/7 mice (Fig. 6d). In the trials in which mice managed to initiate the movements, the reaching delay after the appearance of water droplets was significantly longer. Mice often performed several unsuccessful attempts before a successful reaching were launched (Fig. 6d, Supplementary Video 4). Across all sessions, the movement onset was delayed by 344.7 +/- 84.8 ms (more than 200 ms in 13/20 remaining sessions of 7/7 mice). In line with previous studies ^83–85^, these effects on lift probability and onset timing occurred following targeting of C-C terminals in the IntP, but not the FN (Extended Data Fig. 9).

To further evaluate the role of the C-C projection on ipsi- and contralateral forelimbs, we unilaterally inhibited the C-C terminals in CN (Fig. 6e,f). Photo-inhibition of the C-C terminals ipsilateral to the reaching limb significantly disturbed reaching initiation and/or delayed reach onset, while the effects contralateral to the reaching forearm remain mild at best (Fig. 6e,f). We next analysed the reaching trajectory (Fig. 6g), the endpoint precision (Fig. 6h), as well as the reaching velocity (Fig. 6i,j) in the trials when animals managed to initiate a successful reaching. Interestingly, in contrast to the effects by inhibiting the ponto-cerebellar pathway ^83–85^, inhibiting C-C pathway did not affect the movement kinematics. These results suggest that C-C pathway preferentially affects the timing and sequencing of volitional forelimb reaching, which is distinct from the role of ponto-cerebellar pathways in real-time control of reaching kinematics.

### Distinct task representations of C-C and CSN neurons during volitional forelimb reaching

We have shown that C-C neurons are a subgroup of CSN neurons and they project to a wide range of downstream regions. It is therefore puzzling how the activity of C-C neurons, as a subset of CSNs, does not interfere with the motor commands that control the contralateral limbs during reaching. To disambiguate the contribution of the CSN and C-C neurons in motor control, we investigated their activities during reaching with the contralateral forelimb (Fig. 7a). We injected AAVretro-Cre in PN or CN and AAV-flex-GCamP7f in MC contralateral to the reaching limb to express GCamP7f selectively in CSN and C-C neurons (Fig. 7a). We performed two-photon Ca^2+^ imaging to monitor the activity of the primary dendrites (Fig. 7b). The activities of these dendritic compartments are highly correlated with somatic action potentials ^68,86,87^. The imaging regions for C-C and CSN populations were equally distributed along the anterior-posterior axes of the MC (Extended Data Fig. 10e). We identified all hit, miss and no attempt trials via motion tracking (Extended Data Fig. 10a-d). After the initiation of the first reach, mice continued to retrieve water with multiple sequential reaches. This allowed us to analyse tuning patterns during consecutive sequential reaching movements. Well-trained mice (∼7 days, ∼112.0 ± 3.1 trials/day) almost always lifted the forelimb during the task and successfully retrieved water in over 60% of the trials (Extended Data Fig. 10a-d).

**Figure 7.**
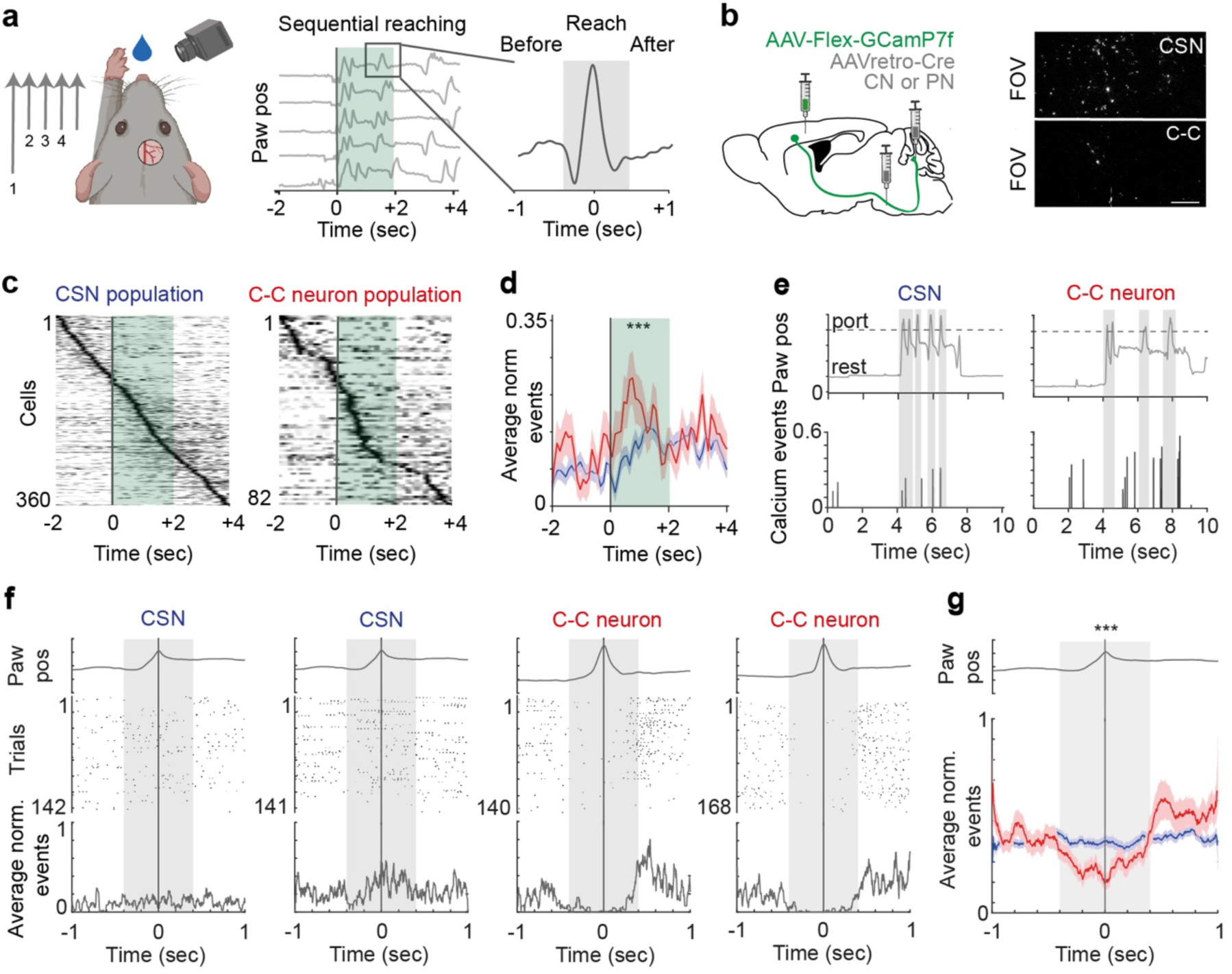
Distinct task representations of C-C and CSN neurons during volitional reaching. a. Schematic of behavioral paradigm. Mice voluntarily reach for a water droplet. After the initial reach, they continue with several sequential reaches towards the water port. The green shading indicates the 2 seconds time window within which the initial reach is executed. The zoom in shows the trajectory of a single reach (gray shading) after the initial reach. b. Viral labelling strategy to image CSNs and C-C neurons contralateral to the reaching arm. Field of view (FOV) shows two-photon images of GCamP7f expression primary dendrites of C-C neurons and CSNs. c. Ca^2+^ events from all C-C neurons and CSNs, aligned to reaching onset, sorted by the timing of their peak modulation. d. Average reaching related activity in C-C neurons and CSNs. The peak modulation in C-C neurons (0.273 ± 0.005 a.u.) precedes the peak modulation in CSNs (0.207 ± 0.005 a.u.) by 500 ms. Values represent the mean ± sem. Two-sample Kolmogorov-Smirnov test, p <0.001. e. Examples of C-C neuron and CSN activity patterns during sequential forelimb reaching. f. Peristimulus histograms for two example CSNs and C-C neurons, showing the activity patterns during sequential reaching epochs when aligned to the time of water port touch. Grey bar indicates 800 ms time window aligned to the time of water port touch. g. Average activity of C-C neurons and CSNs during sequential reaching epochs when aligned to the time of water port touch. Two-sample Kolmogorov-Smirnov test, p<0.0001.

To identify reaching related activity of each cell, we aligned its activity to the onset of the first reach and computed the peri-reaching histogram for each neuron. Subsequent averaging and normalisation revealed the reaching related neuronal activities. In total we recorded the activity of 1444 CSN and 218 C-C neurons; of which 24.9 % (n = 360) CSNs and 37.6 % (n = 82) C-C neurons showed task related modulation, respectively. We sorted all active neurons based on their average peak event timing (Fig. 7c and Extended Data Fig. 10f). While the peak modulations of different CSN neurons were evenly distributed across the entire time window, the C-C population representation was more concentrated within a time window during movement (Fig. 7c). On average, the peak modulation of C-C neurons (0.27 ± 0.01 a.u.) during reaching was significantly higher than that of the CSN population (Fig. 7d, 0.21 ± 0.01 a.u.). During no attempt trials, the activities of both populations remained at baseline levels (Extended Data Fig. 10g), further confirming that these activities are movement specific.

To examine the specific timing of CSN and C-C activities during reaching, we aligned the activity patterns of C-C and CSN neurons to each individual reach (Fig. 7e,f). This analysis revealed that the task representation of CSN neurons is highly variable from trial to trial and equally distributed across the entire reach epoch. In contrast, the C-C neurons were preferentially activated right before and after the reaching window, but were significantly suppressed during the reaching movements (Fig. 7e-g). These data indicate that CSN and C-C neurons encode distinct movement features. The CSN neurons sample broadly all movements features and have highly heterogeneous activity patterns. In contrast, the C-C neurons in the same MC regions preferentially encode information related to the transition of sequential movements. Therefore, the CSN and C-C neurons are likely to be activated in sequence to achieve a fluent control of the movements.

## Discussion

In this study, we identified and characterized a direct cortical-to-cerebellar pathway that establishes direct monosynaptic inputs from the MC to ipsilateral CN, bypassing the conventional multi-synaptic pathways. This direct C-C projection, primarily targeting the FN and IntP, represents a distinct subset of corticospinal projecting neurons. Anatomically, C-C neurons exhibit unique projection patterns and receive predominantly local motor cortical inputs, distinguishing them from traditional CSNs, which have broader cortical inputs and widespread cerebellar innervation via pontine nuclei. Using *in vivo* imaging and optogenetic manipulations, we found that C-C neurons modulate their activity in relation to voluntary movements, especially at movement initiation and during transitions between motor states. Silencing these neurons disrupts ipsilateral limb coordination implicated in both locomotion and skilled forelimb reaching. These findings suggest that the direct C-C pathway may act in concert with the canonical cortico-cerebellar pathways to ensure smooth, bilaterally coordinated motor execution.

### Distinction of the C-C pathway from multisynaptic cerebro-cerebellar systems

Past work has shown that MC and cerebellum interact via elaborate cortico-cerebellar loops, in which the cortex projects to the pontine nuclei, which relays a dense mossy fiber projection to the cerebellar cortex and nuclei ^28–31,80^. This canonical cerebro-ponto-cerebellar circuit is reciprocal: the cerebellar nuclei project back to the thalamus, which in turn influences cortical activity ^12,85^. Within this closed-loop organization, most cerebellar targets lie contralateral to the MC, with the cortical output dominating contralateral control of the body.

In contrast, the C-C pathway provides a monosynaptic shortcut connecting the two major output neurons of the cerebro-cerebellar system: layer 5 CSNs and CN neurons. Although some C-C neurons send collaterals to the pontine nuclei, they co-innervate FN and IntP in ipsilateral CN, a pattern strikingly different from the mostly contralateral innervation of the cerebellar cortex by the cortico-ponto-cerebellar route ^30,31,65^. This ipsilateral projection pattern of the C-C pathway provides a novel pathway that opens the contralateral closed-loop organization of conventional cortico-cerebellar feedback. Functional, it may provide an efficient shortcut with which cerebellar output neurons can receive a copy of MC output, while the majority of motor control signals circulate within the contralateral loop. Further, our rabies tracing experiments revealed that C-C neurons receive proportionally greater input from local motor cortical networks, while the CSNs projecting to pontine nuclei sample widely from sensory and higher-order cortical regions. Thus, at the level of both inputs and outputs, the C-C pathway remains anatomically and functionally distinct from the more diffuse multisynaptic system.

Optogenetic activation of C-C neuron terminals in the CN does not elicit robust, time-locked limb movements in resting animals, implying that this pathway does not drive overt motor output in isolation. Instead, its activity ramps up during behavioral state changes — such as the transition from rest to locomotion or from one reach to the next. Indeed, our imaging experiments revealed that C-C neurons exhibit strong task-related modulation at movement onset and during intervals separating repeated reaches. The preferential engagement of C-C neurons during discrete movement transitions aligns with their limited fine-scale control over movement execution and points to a role in coordinating readiness states and ‘gating’ or ‘switching’ between movement phases. This tuning pattern distinguishes their function from the cortico-ponto-cerebellar efference copy system that transmits continuous kinematic commands from motor cortex to the cerebellum^32,80^.

Behavioral perturbations further support this distinction. While silencing pontine or mossy fiber inputs typically impairs endpoint precision or fine-tuning of forelimb movements ^80,84^, we found that disrupting the C-C pathway alters movement initiation and timing during both locomotion and forearm reaching. The functional distinction of both pathways aligns well with the anatomically distinct innervation pattern in the cerebellum. Anticipatory control and finetuning of movements have been shown to be mediated by high-dimensional computations in the cerebellar cortex ^84^, a region spared by the direct cortico-cerebellar connection but densely innervated by mossy fiber collaterals of the pontine pathway (current study, ^30,31^).

### Unique contribution of the C-C pathway to coordinated movements

It is a well-known principle that MC corticospinal projections are predominantly contralateral, and cortical lesion/stimulation resulted mainly contralateral effects ^12,39,41–43,75,88^. However, ipsilateral representations in MC have also been documented during bimanual tasks or ipsilateral reach movements ^48,49,51,53,54^. How and why the cortex maintains ipsilateral signals in MC has remained an open question ^53^.

Typical CSNs have highly diverse tuning profiles reflecting ongoing kinematics and efference copies of motor cortical command signals ^13,32,44,68,76,89,90^. Our findings suggest that signals transmitted via the ipsilateral C-C pathway complements the motor control signal from the other CSNs. Silencing ipsilateral C-C terminals in CN disrupted motor programs requiring bilateral coordination: mice slowed or halted their locomotion, showed prolonged stance phases, and exhibited asymmetrical gait phases on the balance beam. These perturbations indicate that ipsilateral signals are crucial for properly timed hindlimb movements, consistent with known roles of IntP and FN in postural control and interlimb coordination ^47,70,73^. Accordingly, optogenetic silencing on the side ipsilateral to the reaching forelimb significantly delayed or prevented movement initiation, whereas contralateral silencing produced mild or negligible effects. These findings imply that the direct C-C projection specifically supports movement state-dependent activation of ipsilateral CN outputs. In contrast, the canonical cortico-spinal and cortico-ponto-cerebellar outputs appear to control motor commands and ongoing kinematics via a contralateral dominant route. These different routes might be important for synergistical coordination of different motor commands in a particular sequence.

### CN neurons integrate multiple cortico-cerebellar pathways and modulate movements

The CN are interconnected with various subcortical premotor centres, including the motor thalamus, red nuclei, reticular nuclei, and spinal cord, providing multiple pathways through which the cerebellum can influence movement initiation ^45,85,91,92^. Notably, cerebello-spinal neurons from IntP that project to ipsilateral spinal cord are required for skilled forelimb movements, while projections from IntP and FN to the contralateral red nucleus link the CN with the rubrospinal tract and are essential for locomotive behaviour. The rubrospinal tract then crosses the midline again to innervate the spinal cord on the same side as the original cerebellar nucleus and to coordinate movements on the ipsilateral side of the body ^45,46,91,92^. Both the corticospinal and rubrospinal pathways are crucial for movement execution, working together to coordinate flexor and extensor muscle activity in the fore- and hindlimbs ^93^.

The unique anatomical organization of C-C neurons positions the C-C pathway at the intersection of these two descending motor systems. Specifically, C-C neurons send contralateral projections to the spine while also projecting to the ipsilateral cerebellum, which implies that C-C neurons can simultaneously influence the rubrospinal and corticospinal pathways. By bridging these pathways, C-C neurons might thus help to coordinate ipsi- and contralateral limb movements thereby facilitating bimanual motor control required for the transition between movements. An important future direction will be to dissect precisely how C-C signals are integrated by specific subpopulations of CN neurons, particularly within FN and IntP, and how they interact with the broader cortico-cerebellar loops to optimize bilateral limb coordination and postural stability. Elucidating these mechanisms states could open new avenues for understanding cerebellar contributions to voluntary motor control.

Finally, direct cortico-cerebellar connectivity is not unique to rodents: comparable anatomical pathways have been described in birds, cats, and nonhuman primates^94–97^. Although the functional roles of the cortico-cerebellar projection in other species have not been examined, the prominent ipsilateral projections in multiple species suggests an evolutionarily well-conserved mechanism to coordinate, synchronize and/or sequence bilateral limb movements and stabilize motor transitions. Consequently, the basic organizational and functional principles of the C-C pathway demonstrated here may extend across vertebrates.

## Methods

### Animals

All experiments were performed in accordance with the European Communities Council Directive. All animal protocols were approved by the Dutch national experimental animal committee (DEC). Wild-type C57BL/6J (No. 000664) and transgenic Ai32 (No. 024109) mice were purchased from the Jackson Laboratory. The genotype was tested by PCR reaction using toe-tissue gathered at postnatal day 7-10. The mice used in this study were 6-30 weeks old and individually housed in a 12-hours light/dark cycle with ad libitum access to water and food.

### Viral vectors

For the viral labeling of cerebro-cerebellar projecting cells we used the following viral vectors: retro-AAV2-CAG-Cre, AAV1-CMV-Cre-GFP, retro-AAV2-tdTomato, AAV1-Flex-Tdtomato, AAV9-Flex-GFP. In addition, we used Choleratoxin B (CTB, 1:10000) as a retrograde tracer. For the rabies tracing we combined the retro-Cre injection strategy with co-injection of AAV9-FLEX-H2B-GFP-2A-oG and AAV-Ef1A-DIO-HTB as well as co-injection of Rabies-deletedG-CMV-EnvA-mCherry and CTB. For Calcium imaging and photo manipulation experiments we combined the retro-Cre strategy with AAV9-CAG-FLEX-GCamp7f, AAV1-hsyn-FLEX-ChR2(H132)-tdTomato and AAV1-hsyn-DIO-FLEX-OPN3-mScarlet, respectively. A summary of all injection strategies can be found in Table S1.

### Surgeries

For all surgeries, mice were anaesthetized with isoflurane (5% in 0.5 L/min O_2_ during the induction and 1.5% in 0.5 L/min O_2_ for maintenance). The skull of all mice was fixed on a stereotaxic surgical plate (David Kopf Instruments), the body temperature was maintained at 37 °C and the eyes were covered with Dura Tears (Alcon Laboratories). For the local pain treatment, we applied Lidocaine (2.5 mg/ml) to the skull. Buprenorphine (50 µg/kg bodyweight) was applied at the beginning of the surgery for analgesia.

### Stereotaxic delivery of viral vectors

After the removal of the hair on the scalp, the skin above the skull was opened. Subsequently, bregma and lambda were levelled and small craniotomies were established above the injection sites. For all injections, a glass capillary was filled with viral vectors and lowered to the corresponding coordinates in the mouse brain. A summary of all injection strategies, including viruses, injection volumes and coordinates can be found in Supplementary Table S1. Depending on the experimental strategy, the brains were injected uni- or bilaterally. After surgery, the skin was sutured and the animals recovered for at least 5 days.

### Craniotomy

After incubation for 6-8 weeks, a craniotomy of approximately 3 mm length was placed above the FN and IntP for in-vivo recordings. For constructing the recording chamber, a thin layer of Optibond All-in one (Kerr) was distributed on the surrounding skull. The chamber was built with the use of Charisma (Heraeus Kulzer) and sealed with Picodent twinsil after surgery and recording.

### Cranial window implantation

The cranial window was built by attaching a round cover glass (4 mm in diameter) to a custom-made metal ring of 0.5 mm height. The surface of the skull was covered with a thin layer of Super-Bond activator (Superbond C&B, Sun Medical Co., Furutaka-cho, Japan). A circular craniotomy (4 mm in diameter) was established, with its centre 1 mm anterior and lateral to bregma (unless otherwise stated in the figure legend). After viral injection of retrogradely transported cre to PN or CN and cre-dependent GCamP7f to MC, the cranial window was implanted above the brain surface and glued to the brain. Dental cement (Superbond C&B, Sun Medical Co., Furutaka-cho, Japan) was used to fix a metal bar on the skull surface surrounding the cranial window.

### Optic fiber implantation

For optogenetic stimulation, 3.5 mm long optical fibres (200 µm core, 0.22 NA, Thorlabs) were inserted through a craniotomy. For bilateral stimulation one fibre was implanted without ancle above CN (in mm: -1.3 (x), -2.6 (y), -2.3 (z)) and one fibre was implanted with a 15 degrees angle (in mm: -1.6 (x), -2.6 (y), -2.4 (z)). Both fibres were chronically fixed to the skull with Charisma (Kulzer, Hanau, Germany) and Optibond (Kerr Dental, Herzogenrath, Germany). The head bar was fixed with an additional layer of dental cement.

### Reaching Behaviour

Mice were trained to reach for water under head-fixed condition ^75^. In this behavioural reaching paradigm, a tone was followed by a 2 seconds delay period and the appearance of a water droplet. Mice had to learn to detect the position of the port and to reach for the water drop after the delay period. The behavioural paradigm was controlled by a Bpod State Machine (San works LLC, Rochester, United States), which interfaced with water valves, speakers, infrared video cameras and pulse generator for the light stimulation. The reaching behaviour was videotaped with an infrared USB camera (BASLER acA640 – 750 µm, Basler AG, Ahrensburg) at 250 frames/second. An infrared right-angle prism (MRA25-E02, Thorlabs) was positioned below the water port to provide a bottom view of the movement. Two weeks after surgery, the access to water was restricted and mice received 1 ml of water per day until they reached a minimum of 80% of their initial body weight. During the initial water-deprivation period, they were habituated to the head-fixation in the mouse holder and the experimental setup. Once fully water-deprived, mice were trained to reach for a 5 µl water drop that was placed 5 mm below the tip of their nose. Initially the whisker pad was tickled during the presentation of the water drop, which induced reflexive grooming and allowed the mouse to transition to voluntary forearm reaching. After 1-5 training sessions, mice have learned to voluntarily reach for the water and to adjust to the 2 seconds delay period. The inter-trial interval was randomized to 7-14 seconds, and reaching movements were recorded for up to 300 trials.

### Photo-inhibition during Reaching Behaviour

Optogenetic silencing was performed by optical stimulation of OPN3 expressing axonal fibres in the CN. For these experiments, we recorded 50 reach trials before and 30 reaches after optogenetic silencing. Bpod behavioural system and Pulse Pal (Sanworks LLC, Rochester, United States) in combination with a high-power light driver (DC2100, Thorlabs) and LED at 530 nm, were used to deliver 5 minutes of 50 Hz light pulses at 2 mW of light at the tip of the fibre.

### Behavioural Analysis of Reaching Behaviour

For behavioural analysis, Deep Lab Cut software was used to extract the x-, y- and z-coordinates of mouse forearm position during reaching movements (Mathis *et al.*, 2018; Nath *et al.*, 2019). Custom written analysis scripts were used to classify behavioural videotapes into hit, miss and no attempt trials. The performance rate is defined by hits / all trials; lift probability is determined by hits plus misses/all trials and first hit probability is calculated by hits/hits plus misses. Sessions with performance rate below 50% were excluded from analysis. For the analysis of movement kinematics, sessions with more than 5 remaining reaches were analysed and all hit trials were analysed for reaching onset as well as reach velocity and endpoint. The timing of movement onset was defined by the disposition of the reaching forearm by 5 mm from the resting position of the bar. Movement velocity was calculated by the frame-to-frame spatial disposition of the forearm in the x-direction. Movement endpoints were defined by the x-, y- and z-corresponding values of maximal forearm disposition relative to the water port. For the plotting of the average 3D-trajectory as well as average velocity, all ‘hit’ trials were aligned to movement onset and averaged across trials.

### Two-Photon Calcium Imaging during Reaching Behaviour

To study the activity of cerebro-cerebellar projecting cells during forearm reaching movements, well trained GCamP7f expressing mice were imaged in a two-photon microscope (A1RMP, Nikon) that was controlled by Nikon imaging software (NIS Elements, Nikon). We used Bpod State Machine (Sanworks, US) interface to control Nikon imaging software, infrared camera videotaping, water valves and speakers. To synchronize behavioural videotaping and two-photon image recording, individual frame acquisition of both systems was sampled to a NIDAQ-board (National Instruments, Austin, USA). In the imaging system, a 25X objective (Nikon) and a resonant scanner system (A1-SHRM, Nikon) were used to image a 512-by-512 µm FOV at 15.2 frames/second. For excitation a Ti:Sapphire ultrafast pulsed laser (Mai Tai HP, Spectra-Physics) at wavelengths of 920 nm for GCamP7f and 960 nm for RFP at a maximum of 70 mW was used. Images were acquired by a photomultiplier tube. Per imaging session, 90-150 trials were sampled with a randomized inter-trial-interval of 20-30 seconds. In each trial, a 2 second baseline, 2 second delay between tone and water and a 8 seconds response epoch were recorded. Imaging depth was set to 300 µm below the surface, which allowed monitoring of calcium events in tufted dendrites of L5 neurons. The imaging FOV was located by using stereotactic coordinates and distributed across the full anterior-posterior axis of the cranial window (-1 mm to + 3 mm from Bregma). In a small proportion of CSN and C-C neurons we applied widefield calcium imaging at a sampling rate of 30 frames/second. Both two-photon and wide-field datasets were merged.

### Two-Photon Image Analysis

Calcium imaging analysis was performed using CNMF ^86^. Imaging datasets lasting 90-150 trials (12 seconds per trial) were merged and movement corrected using rigid, followed by non-rigid registration ^86^. Movement-corrected datasets were then analysed using CNMF, where dendrites with temporally and spatially overlapping activity patterns were identified to originate from the same cell and thus excluded from analysis. The output from this analysis delivered ΔF/F signals and their deconvolved events. Subsequently, ΔF/F signals and extracted events were aligned with the x-, y- and z-coordinates of the reaching forearm, previously extracted with Deep Lab Cut based video-tracking algorithms (see above). All trials were classified into hit-, miss- or no-attempt trials. For movement-related signal detection all hit trials were aligned to movement onset. Then tone as well as movement-aligned events across all hit trials were multiplied by calcium amplitude and averaged. We applied a threshold of 10 events across all trials to identify all neurons active during the execution of the task. The event rates of each cell were normalized to the maximum and sorted by peak-event timing. Tone and movement-related modulation patterns were detected by the timing of the peak event rate during time windows corresponding to tone (within 1 second after the presentation of tone) or movement (2 seconds after movement initiation). For the analysis of calcium activity during sequential reaches, time windows of 1 second before and after each touch were aligned, averaged and analyzed.

### Treadmill

In this behavioural paradigm, mice were trained to walk on a treadmill under head-fixed conditions, without any stimuli, to study differential cell activity in running or resting states. During 7 consecutive days, mice were trained for 60 minutes in the setup. Training sessions included artificial movement of the wheel when mice were at rest for long periods of time to spur their movement. In the following days, mice were set on the treadmill for 5 minutes before the start of recordings. Behaviour recordings were done with a USB camera (BASLER acA640 – 750 µm, Basler AG, Ahrensburg) at 250 frames/second, while widefield recordings were done with a Kinetix sCMOS camera (Teledyne Photometrics), at 30 frames/second. Both cameras were synchronized by a Bpod State Machine (Sanworks LLC, Rochester, United States).

### Widefield Imaging for Treadmill

Well-trained GCamP8f-expressing mice were imaged in the one-photon setting of a multiphoton microscope (A1RMP, Nikon). In the imaging system, a 25x objective (Nikon) was used to image a 832x832 µm FOV. For excitation a LED light was filtered by 482/35 filter cube (SemRock Brightline FITC-3540C) at powers lower than 6mW (7.2mW/mm^2^). Images were acquired with a Kinetix sCMOS camera (Teledyne Photometrics), controlled by MicroManager. All locations with active cells were recorded for 5 minutes, and their locations were labelled based on stereotactic coordinates. Mice were kept on the setup for recordings for a maximum of 1.5h hours per day, and only recordings with at least 1 minute of walking and 1 minute of resting were accounted for analysis.

### Behavioural Analysis for Treadmill

For behavioural analysis, Deep Lab Cut software was used to extract left and right paw coordinates during movement ^81,82^. Customized analysis codes were used to classify into resting or running states. Movement traces were filtered by a 1ms rolling median to remove outliers and smoothened with a 5 ms Savitszy-Golay filter. Speed was calculated as the average distance between 2 time points of both paws. Acceleration was calculated as speed over time. Running was classified as at least 3 seconds of continuous movement. Movement start was defined as the start of a 3 seconds uninterrupted period of movement, and movement stop was defined as the beginning of 3 seconds of continuous movement interruption. For visualization purposes, the speed and acceleration traces were binned to match the widefield recording frequency.

### Widefield Image Analysis

Imaging datasets were movement-corrected using rigid registration ^86^ and then analyzed with OnACID, from which we obtained the deconvolved calcium traces. We detected neurons on an initial batch of 3000 frames and further deconvolved the remaining frames. For all analysis, we used the C variable (deconvolved trace) of the OnACID output. For post-processing, we started by excluding cells with less than 10 spikes over the 5-minutes recordings. Then, we classified each cell as movement preference cell (MPC), resting preference cell (RPC) or a non-preferential cell (NPC). MPCs and RPCs were defined as having at least 85% of their spikes in either movement or resting epochs, respectively. Decelerating and Acceleration cells were classified as cells that had an average spike peak correlated with the lower or higher 10% of their acceleration, respectively.

### Balance Beam

The balance beam behavioural paradigm was used to study balance in mice. A metal beam with a diameter of 14 mm in diameter was placed between two stages lifted 20 cm above ground. While mouse walked across the beam, we applied a 50 Hz photo stimulus for 5 second sand monitored the body position with a GoPro (HERO8 Black) from the back of the mouse. For photo-inhibition we applied 5 seconds 50 Hz pulse train (at 530 nm and 2 mW). For photo-activation, we applied a 50 Hz photo stimulus for 5 seconds at 470 nm. Behavioural analysis was performed with Behavioural Observation Research Interactive Software (Boris) a behavioural event logging software for video coding and live observations (Friard & Gamba, 2016). Behavioural events such as slipping with the left or right hind paw, freezing, stopping and balance adjustments have been ranked. For the control condition, 5 second walking equivalent to the photo-stimulation period were analysed.

### Locomouse and Photo-inhibition

A custom-designed setup was developed to assess whole-body coordination during locomotion in mice. The Locomouse setup consists of a glass corridor under which a 45-degree mirror is placed to provide a bottom view of mouse locomotion. Mice were videotaped crossing the monitor with a high-speed infrared USB camera (BASLER acA640 – 750 µm, Basler AG, Ahrensburg) at 400 fps and a spatial resolution of 1400 x 230 pixels. After initial habituation, mice walked freely between two boxes at the ends of the glass corridor. While mice were crossing the glass corridor, a 5-second light pulse (50 Hz, 2 mW) at 530 nm was applied, and the effect on walking patterns was monitored.

### Analysis Locomouse

We used DeepLabCut ^81,82^ video tracking software to extract the coordinates of nose, left and right fore-as well as hind-paws and the tail in the side and bottom view of the Locomouse. We used custom written Matlab analysis scripts to calculate the velocity of all four paws, which allowed the analysis of individual stride cycles. Individual strides were subdivided into swing and stance phases, which allowed us to calculate the following parameters:

Stride length: disposition of the forepaw from start of stance to start of next stance in x.

Stride duration: duration of forepaw displacement between two stances.

Swing length: disposition of the forepaw during swing.

Swing velocity: speed of forepaw displacement during swing.

Stance duration: time that the forepaw is placed on the ground before the next swing is initiated.

Stance phase: (stance time – stance time _Reference paw_)/stride duration

### iDISCO+ Brain Clearing Procedure

For the iDISCO clearing we followed the protocol published earlier ^59^. In the first step, the brain was dehydrated by 1hour of incubation steps in methanol solutions (20%, 40%, 60%, 80% and 100%) and overnight in 1/2 mix of methanol dichloromethane. Next, we applied a bleaching step in 5% H_2_O_2_ in 90% methanol at 4°C overnight. After re-hydration, the brain was incubated in blocking solution for 2x at 37°C. Afterwards the brain was incubated for 1 week at 37°C in 4 ml primary antibody solution that contained PTwH incubation medium (PBS, 0.2% Tween-20, 100 µg/10ml heparin, 5% DMSO, 3% Normal Donkey Serum) and anti-GFP antibody (35, Abcam,) in a dilution of 1:1000. The incubation with primary antibody was repeated 2x for a week. After washing, the brain was incubated for 1 week in 4 ml secondary antibody solution that contained anti-rabbit Cy5 (4) diluted in PTwH (1:750). All antibody-containing solutions were filtered at 0.2 µm and amphotericin (10 µl/ml) was added every second day to prevent fungal growth. All antibody incubation steps were performed at 37°C. After washing, the final tissue clearing step was performed at room temperature. The brain was dehydrated by 1hour of incubation steps in methanol solutions (20%, 40%, 60%, 80% and 100%) and overnight in 1/2 mix of methanol/dichloromethane. The following day, the brains were placed dichloromethane (2 times 20 minutes) and final clearing was achieved by storage in benzylether.

### Light-sheet imaging

Whole brains (the cerebellum was partially removed) were imaged on a light-sheet microscope (Ultramicroscope II, LaVision Biotec) with a Neo sCmos camera (Andor, 2560 x 2160 pixels). Samples were imaged with a double-sided illumination and a sheet NA of 0.148348, which results in a 5 µm thick light sheet at step-sizes of 2.5 µm. The effective magnification for all images was 1.36x (zoom body*objective plus dipping lens = 0.63x * 2.152x). For the illumination, we used the Coherent OBIS 488-50 LX Laser with a 525/50 nm filter for the autofluorescent channel and the Coherent OBIS 647-120 LX with the 676/29 nm filter for the fluorescent channel.

### Image Processing and Clearmap Analysis

All cell bodies were manually mapped as spots with the use of Imaris software (Bitplane, http://www.bitplane.com/imaris/imaris). A 3D reconstruction of the location of the cell bodies was used as an input file for the cell detection and registration pipeline ClearMap ^98^. In short, the ClearMap platform registers the brain to a reference atlas, detects the location of cells, maps them to the reference atlas, and generates heatmaps of the cell locations as well as statistical analysis.

### Single-Axon Tracing

For the single-axon tracing experiments, mice were perfused, and their brains were stored in PFA. The histological tissue processing and imaging, as well as the single axon tracing and reconstruction, followed the pipeline published previously ^99^.

### Immunohistochemistry and Image Acquisition

For histology the mice were perfused with 4% paraformaldehyde (PFA, in 0.1 M phosphate buffer (PB) at pH 7.3), post-fixed in PFA for 2 hours and stored in 10% sucrose overnight. After removal of the dura, the brains were embedded in a 12% gelatine and 10% sucrose solution and incubated in 10% formaline and 30% sucrose solution for 3 hours. Afterwards the brains were stored in 30% sucrose. The brains were cut in slices of 50µm thickness on a cryotome (SM2000R, Leica) and stored in 0.1 M PB. Before antibody treatment slices were blocked for 2 hours at RT (10% normal horse serum (NHS) and 0.5% Triton-X100 in PBS). Primary antibodies were incubated overnight at 4°C (2% NHS and 0.5% Triton-X100 in PBS) and secondary antibodies were incubated for 4 hours at RT. Brain slices were rinsed and mounted in vectashield (Vector Laboratories). The following primary antibodies were used for fluorescent stainings: anti-RFP (rabbit, 1:2000, Rockland), anti-CTB (goat, 1:15000, List labs). The following secondary antibodies were used: anti-rabbit-Cy3 (1:400, Jackson Immunoresearch), anti-goat-Cy5 (1:400, Jackson Immunoresearch). For the background staining of cell nuclei, all sections were incubated for with DAPI (300 nM). The fluorescent overview images were acquired with a 10x objective on a fluorescent scanner (Axio Imager 2, Zeiss). The confocal images were acquired on a LSM 700 microscope (Zeiss) and the use of 20X, 0.3 NA as well as 63X, 1.4 NA objectives, respectively.

### Histological Analysis

Histological analysis of axonal innervation in CN was performed with AMaSiNe analysis software ^100^. In short, the fluorescent signal of individual sections was warped to 3D Allen Brain Reference Atlas and the labeling was reconstructed. This allows precise comparison of multiple brain data sets in Allen Brain Space and reveals spatial representation of neural projections.

### Statistics

For statistical analysis, all data are represented as the mean +/- standard error of the mean (SEM). For statistical analysis GraphPad PRISM software package and Matlab were used.

## Author Contributions

C.S. and Z.G. conceived this study and contributed to study design. C.S. coordinated and conducted experiments with the contribution from other authors. R.O.S. contributed to *in vivo* imaging experiments. Y.A. and J.P. contributed to the iDISCO imaging. X.L., A.L., and H.G. contributed to fMOST experiments. C.S., R.O.S., H.S., and H.H. analyzed the data with assistance with from other authors. All authors contributed to the drafting of manuscript with important intellectual content. C.D.Z., F.H. and Z.G. jointly supervised the project.

## Acknowledgement

We thank E. Goedknegt, E. Haasdijk and M. Rutteman for their support with histological analysis. We want to thank H. Schäfer, B. van Hoogstraten, B. Bouwen, R. Vos, M. Cozzolino, M. Berrin Gül, S. Greenthaner and G. Getachew for their help with experimental work. We thank N. Li, J. Zhu, D. Huber, G. Galiñanes, and M. Prsa for comments on the manuscript. This work is supported by NWO VIDI grant (Z.G., VI.Vidi.192.008), NWO-Klein grant (Z.G., OCENW.KLEIN.007) and ERC-stg grant (Z.G., 852869).

## Data availability

The data generated in the current study is available from the corresponding authors on request.

## Extended Data Figures

**Extended Data Figure 1.**
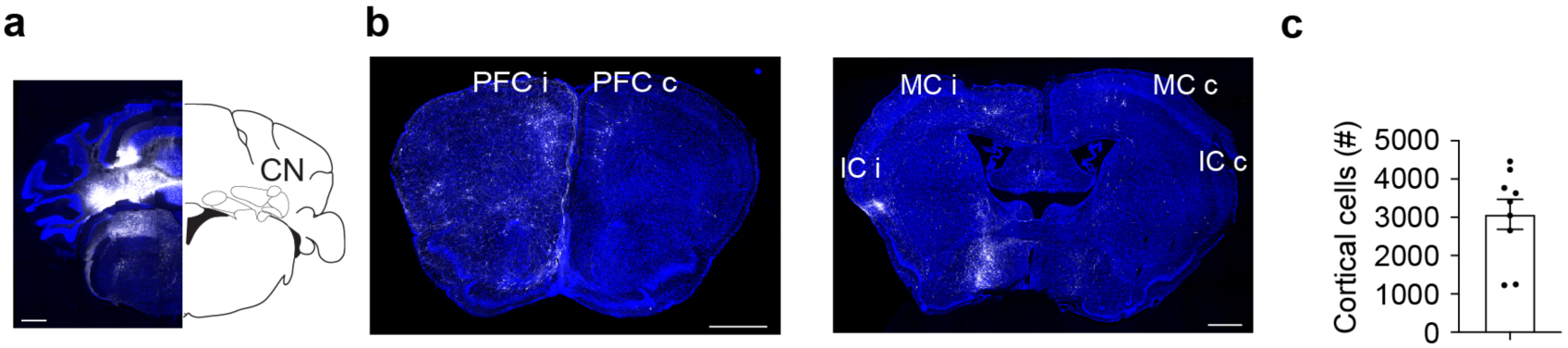
Quantification of cortical neurons that project to CN using retrograde tracing. a. Widefield image showing a unilateral injection cite in the CN. Scale bar 500 µm. b. Transverse sections of retrogradely labeled cells in ipsilateral (i) as well as contralateral (c) prefrontal cortex (PFC), motor cortex (MC) and insular cortex (IC). Scale bar 500 µm. c. Quantification of the total cortical-cerebellar projecting neurons across different animals. Neurons were quantified from 1 in 4 histological sections. N = 9.

**Extended Data Figure 2.**
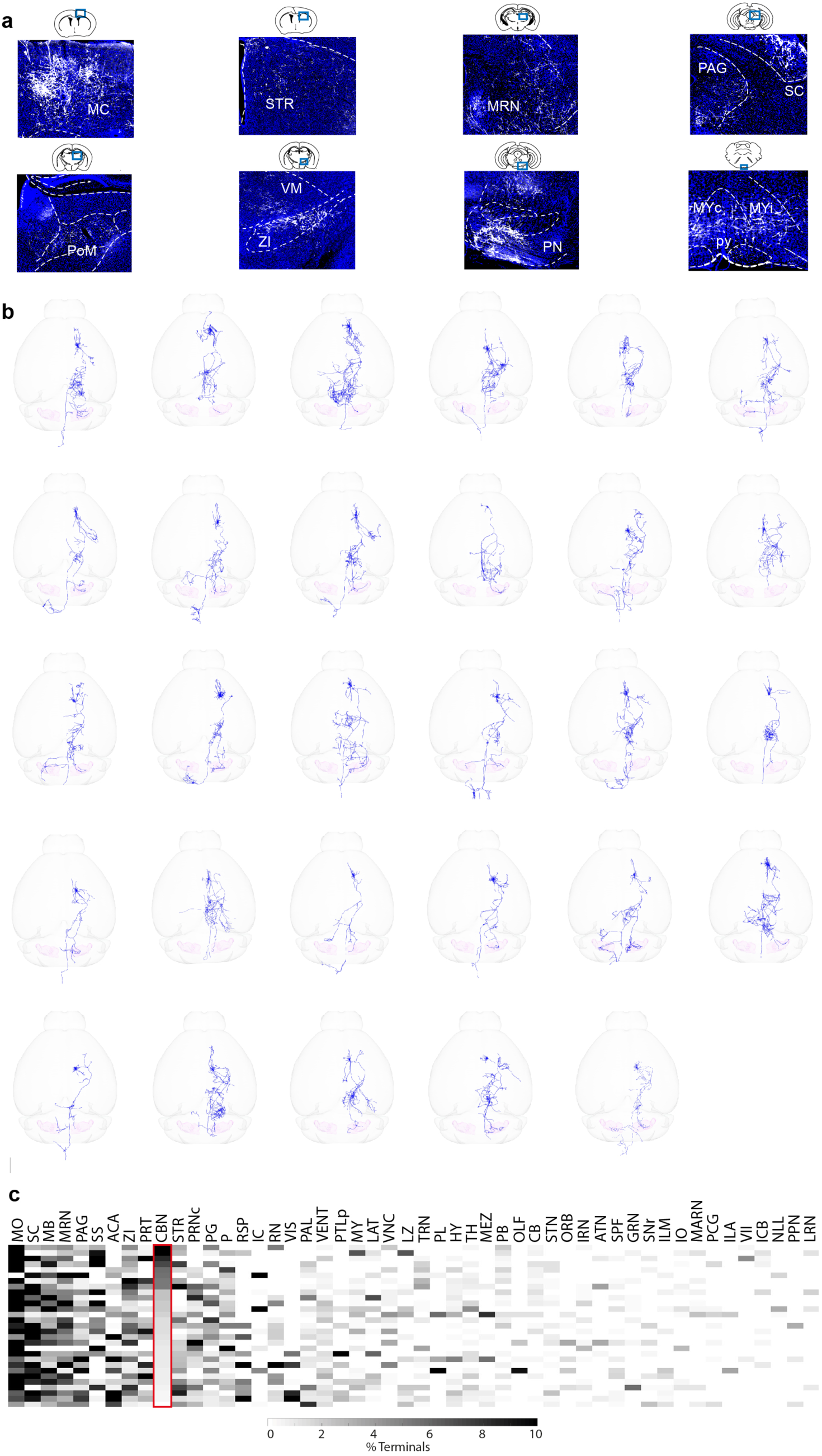
Single axon tracing of C-C neurons in the whole brain. a. Transverse sections depicting C-C neuron collaterals throughout the brain. Regions include Motor cortex (M1), striatum (STR), midbrain reticular nucleus (MRN), periaqueductal gray (PAG), superior colliculus (SC), posterior medial nucleus of thalamus (PoM), ventromedial thalamus (VM), zona incerta (ZI), pontine nucleus (PN), medulla (MY), pyramidal tract (py). Scale bar 200 µm. b. 3D reconstruction and rendering of the projection pattern of individual C-C neurons. c. Heatmap showing the fraction of terminal endpoints of C-C neurons in all relevant downstream regions. Rows represent individual cells that are sorted based on their cerebellar innervation strength. Columns represent target areas.

**Extended Data Figure 3.**
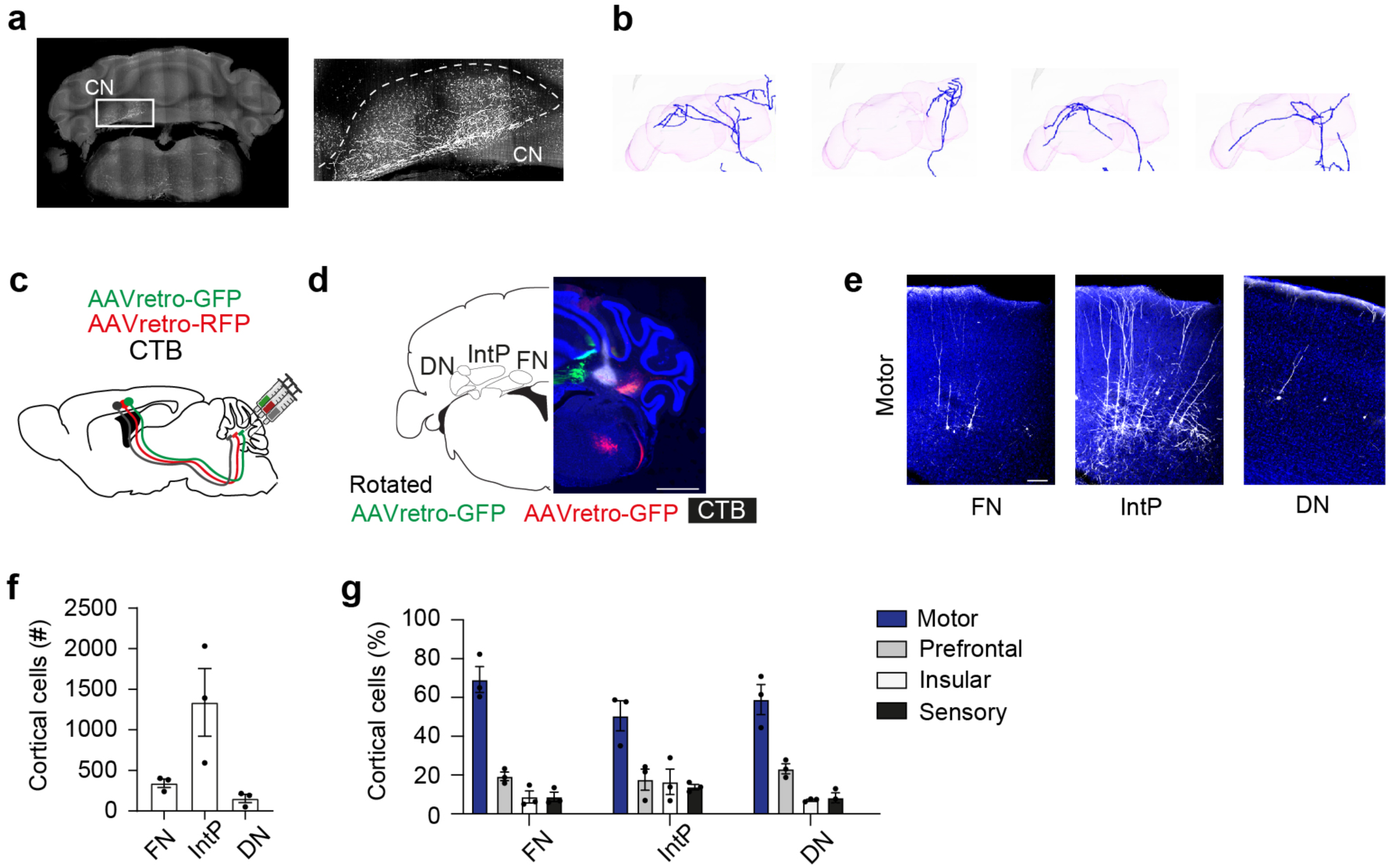
Cerebellar nuclei specific innervation by C-C neurons. a. Image depicting motor cortical innervation in CN after fMOST imaging. b. Single-axon reconstruction of motor cortical terminals in CN from 4 example C-C neurons. c. Retrograde labeling strategy to map nucleus specific inputs from cortex. Three retrograde tracers (AAVretro red-fluorescent protein (RFP), AAVretro green fluorescent protein (GFP) and Cholera-Toxin B (CTB) were delivered to fastigial nucleus (FN), interposed nucleus (IntP) and dentate nucleus (DN). d. Example image of retrograde tracer injections in DN, IntP and FN. Scale bar 1000 µm. e. Example labeled motor cortical neurons that project to FN, IntP and DN respectively. Scale bar 100 µm. f. Quantification of retrogradely labeled neurons in cortex after injection to FN, IntP and DN. g. Quantification of retrogradely labeled neurons in motor, prefrontal, insular and sensory cortex.

**Extended Data Figure 4.**
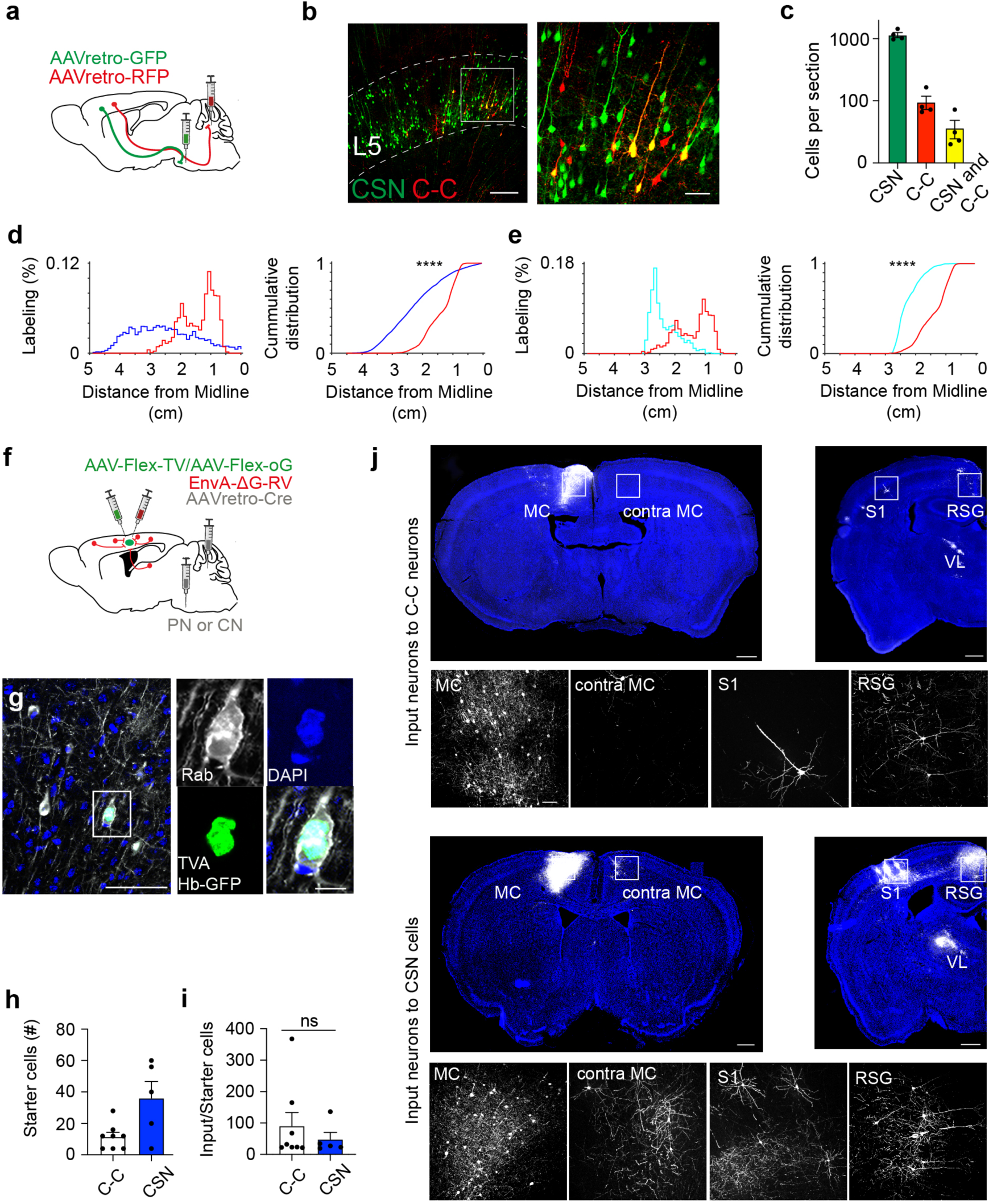
Characterization of the C-C and CSNs pathways. a. Viral strategy to deliver retroAAV red-fluorescent protein (RFP) to CN and AAVretro green fluorescent protein (GFP) to PN. b. Confocal image showing pontine projecting CSNs (in green) and C-C neuron (in red) in the layer 5 of motor cortex. Scale bar 200 µm. Zoom in image showing overlapping labelling in a subpopulation of neurons. Scale bar 50 µm. c. Quantification of cortical neurons that project to PN, CN or both. n = 4, PN: 1149 ± 104.4, CN: 95.75 ± 23.26, PN and CN: 36.5 ± 12.1 cells per section. d. Proportion and cumulative distribution of mossy fiber boutons in cerebellar cortex (blue) and C-C inputs in CN along the mediolateral axis of the cerebellum. Kolmogorov-Smirnov test, p = 0.000631. e. Proportion and cumulative distribution of mossy fiber collatorals in CN (cyan) and C-C inputs in CN along the medio-lateral axis. Kolmogorov-Smirnov test, p = 0.000000007. f. Schematic of rabies labeling strategy fro mapping cortical input neurons to C-C neurons and CSNs. g. Starter cells that co-express TVA-Histone-bound-GFP (TVA Hb-GFP) and Rabies (Rab). Scale bars 50 µm and 10 µm. h. Total number of starter cells for C-C neurons and CSNs. N = 8 and 5, respectively. i. The ratio between the number of input and starter cells for C-C neurons and CSNs. N = 8 and 5, respectively. Mann-Whitney test, p = 0.8329. j. Example coronal sections showing the labeling of the input neurons to C-C neurons and CSNs. The boxed regions depict brain areas with labeling and high magnification images of labeled input cells in local-, long-range cortical and subcortical areas. Scale bar 500 µm and 50 µm.

**Extended Data Figure 5.**
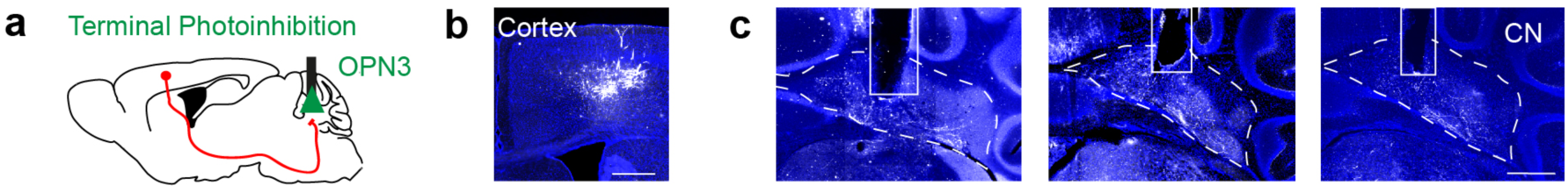
Experimental strategy for photo-inhibition of C-C terminals in CN. a. The strategy for photo-inhibition experiments. b. Image of OPN3 expressing C-C neurons in the MC. Scale bar 200 µm. c. Three example images showing the OPN3 expressing C-C neuron terminals and optic fiber implantation tracks in CN.

**Extended Data Figure 6.**
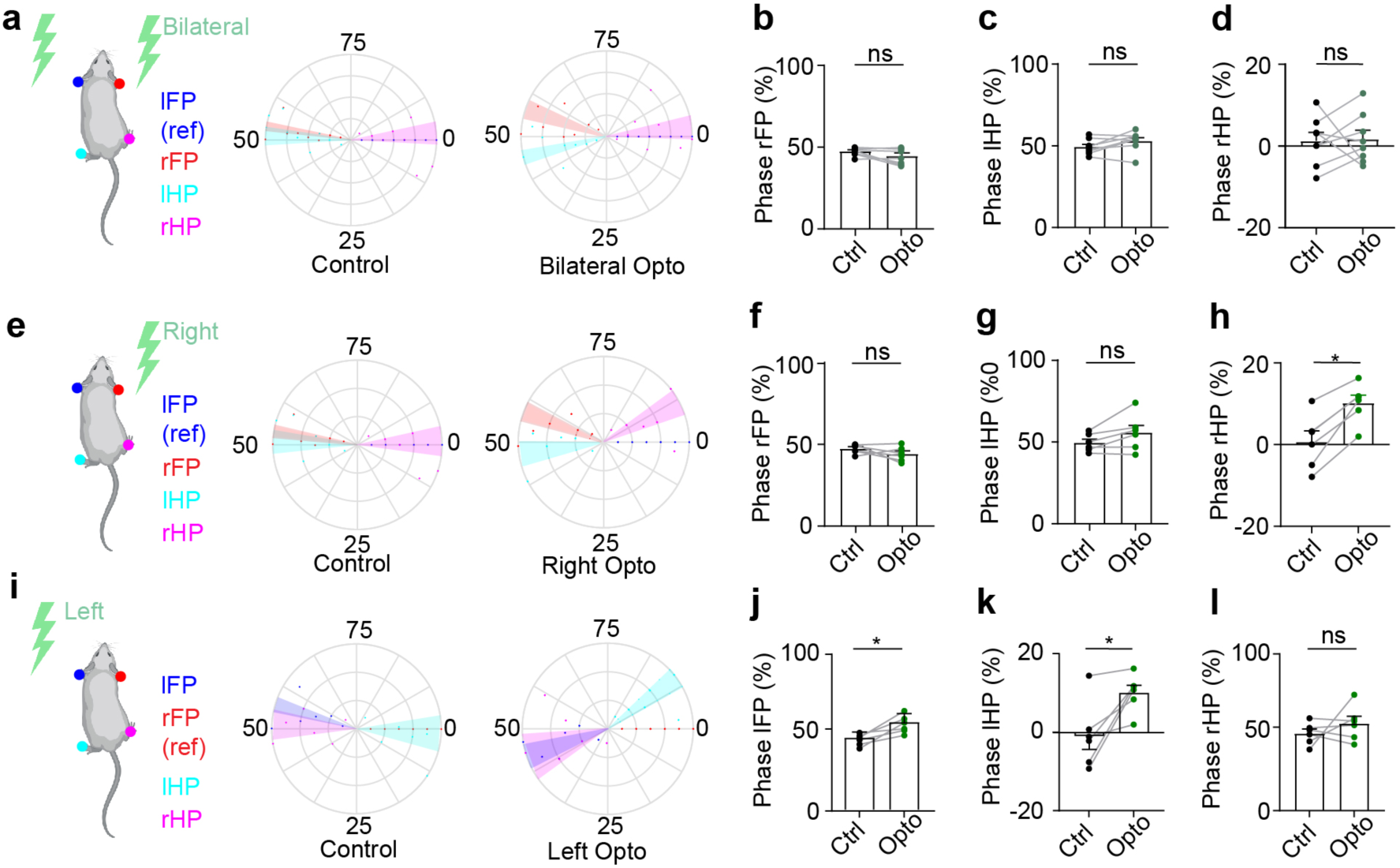
Behavioural analysis of locomotion phases. a. Polar plots indicate the phase of step cycle in which each limb enters stance. Plots are aligned to stance onset of left forepaw for control and bilateral photo-inhibition. Each dot represents one mouse and the shaded area represents s.e.m.. N = 8. Left forepaw (lFP), right forepaw (rFP), left hindpaw (lHP) and right hindpaw (rHP). b. Quantification of the phase of step cycle in which rFP enters stance. N = 8, paired t-test, p = 0.0795. c. Same as (b), but for lHP. N = 8, paired t-test, p = 0.0959 d. Same as (b), but for rHP. N = 8, paired t-test, p = 0.8819. e. Same as (a), but for photo-inhibition in the right CN. N = 6. f. Quantification of the phase of step cycle in which rFP enters stance. N = 6, paired t-test, p = 0.2140. g. Same as (f), but for lHP. N = 6, paired t-test, p = 0.2813. h. Same as (f), but for rHP. N = 6, paired t-test, p = 0.0120. i. Same as (a), but for photo-inhibition in the left CN. The rFP was chosen as reference paw. N = 6. j. Quantification of the phase of step cycle in which lFP enters stance. N = 6, paired t-test, p = 0.0104. k. Same as (j), but for lHP. N = 6, paired t-test, p = 0.0199. l. Same as (j), but for rHP. N = 6, paired t-test, p = 0.3676.

**Extended Data Figure 7.**
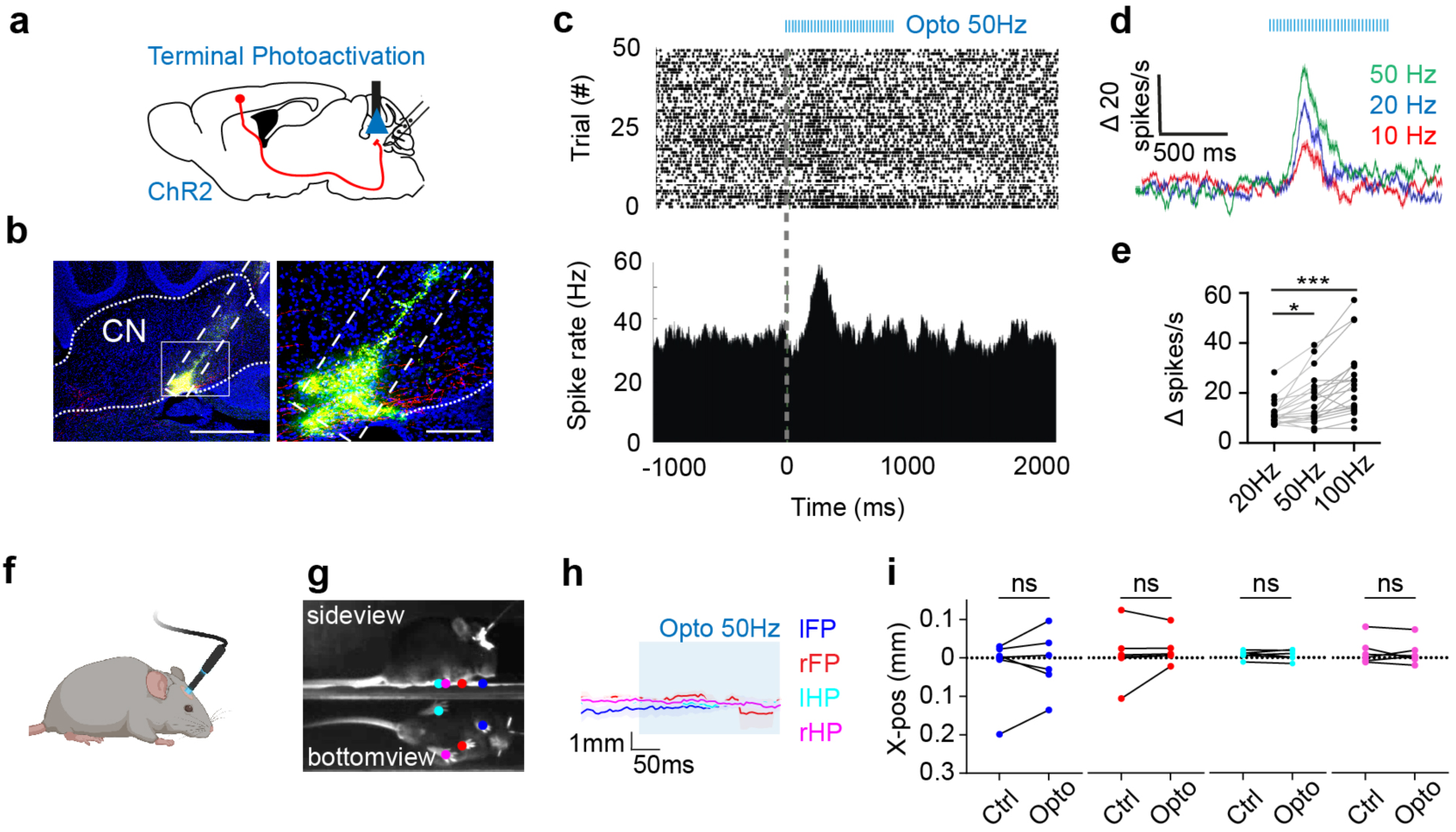
Photo-activation of C-C neuron collaterals facilitates CN neuron activity, but does not induce movement. a. Illustration of viral labeling, photo-stimulation and recording strategy. b. Left: image of a DiO labelled recording track in CN. Right: high magnification image of the recording track and the ChR2 expressing C-C neuron terminals in CN (red). Scale bars 500 µm and 50 µm. c. Peri-stimulus time histogram shows CN neuron response to C-C neuron collateral stimulation. d. The increase in spike rates of all responsive cells at different photo stimulation frequencies. e. Quantification of the average modulation rates in response to opto-stimulation at 20, 50 and 100 Hz. Friedman test corrected for multiple comparisons; 20 Hz vs. 50 Hz: p = 0.0923, 50 Hz vs. 100 Hz: p = 0.0261, 20 Hz vs. 100 Hz: p < 0.0001. f. Schematic showing photo-activation of C-C neuron collaterals in CN in a unrestrained resting mouse. g. DeepLabCut based motion tracking of paw positions during photo-stimulation. The bottom view of the mouse is visualized with a mirror placed 45 degrees below the mouse. h. The positions of all paws (left forepaw (lFP), right forepaw (rFP), left hindpaw (lHP) and right hindpaw (rHP)) before and during photo-stimulation. i. Quantification of the paw position displacement before and during photo-stimulation. N = 6, lFP: p = 0.5226, rFP: p = 0.4779, lHP: p = 0.5834, rHP: p = 0.6714.

**Extended Data Figure 8.**
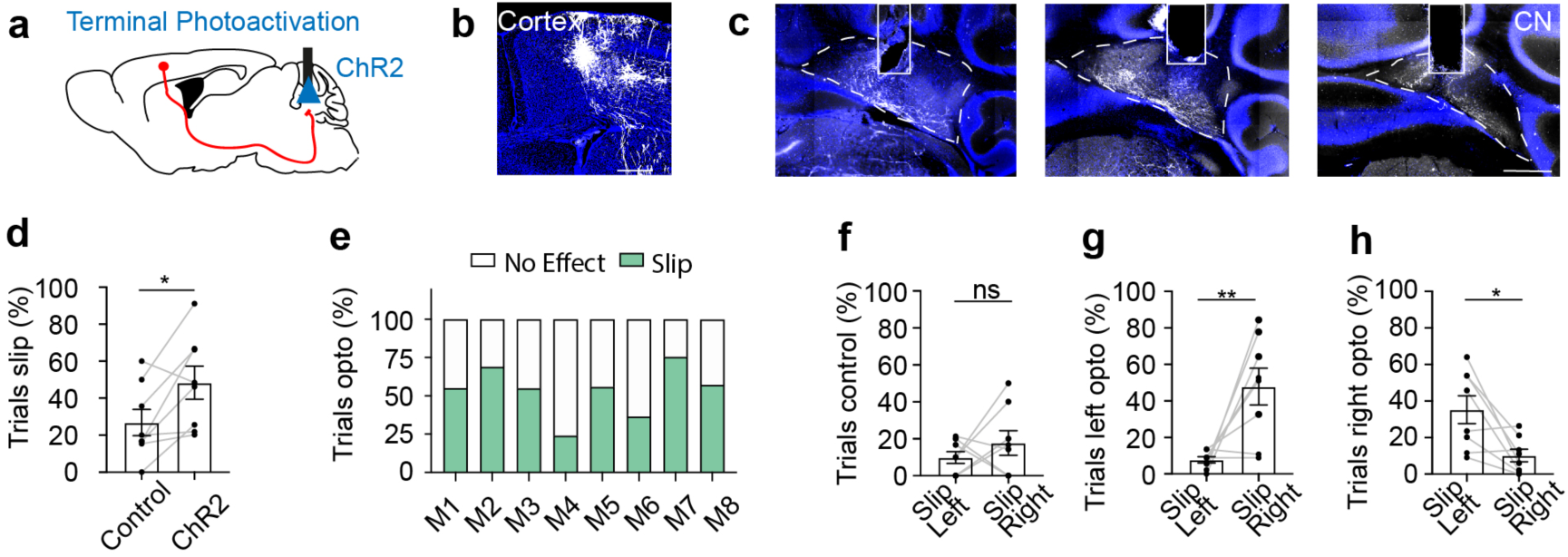
Photo-activation of C-C neurons collaterals induces directional slips on balance beam. a. The viral strategy for photo-activation experiments. b. Image of Channelrhodopsin2 expression in C-C neurons. c. Example CN images from three mice showing the expression of Channelrhodopsin2 in C-C terminals in CN and optic fiber implantation tracks in CN. d. The percentage of trials in which the mouse slips during control and photo-activation conditions. N = 8, paired t-test, p = 0.0236. e. Quantification of the percentage of trials in which the mouse slips after unilateral photo-inhibition. M1-8 indicate individual mice. f. The percentage of trials in which the left or the right hindpaw slip. N = 8, paired t-test, p = 0.3475. g. Same as (f) but for the trials with photo-stimulation of C-C terminals in the left CN. N = 8, paired t-test, p = 0.0100. h. Same as (f) but for the trials with photo-stimulation of C-C terminals in the right CN. N = 8, paired t-test, p = 0.0103.

**Extended Data Figure 9.**
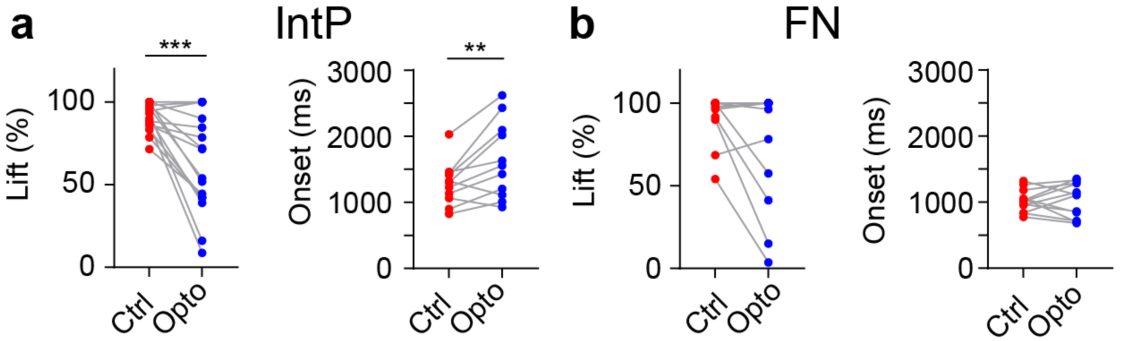
The reaching behavior is affected by photo inhibition of C-C terminals in IntP, but not in FN. a. The lift probability and reach onset timing in all sessions from the animals with fiber implantation in the interposed nucleus (IntP). N = 3, Wilcoxon test, Lift probability: n = 10, p = 0.0004, Onset: n = 11, p = 0.0098. b. The lift probability and reach onset timing in all sessions from the animals with fiber implantation in the fastigial nucleus (FN). N = 4, Wilcoxon test, Lift probability: n = 14, p = 0.2871, Onset: n = 11, p = 0.9697.

**Extended Data Figure 10.**
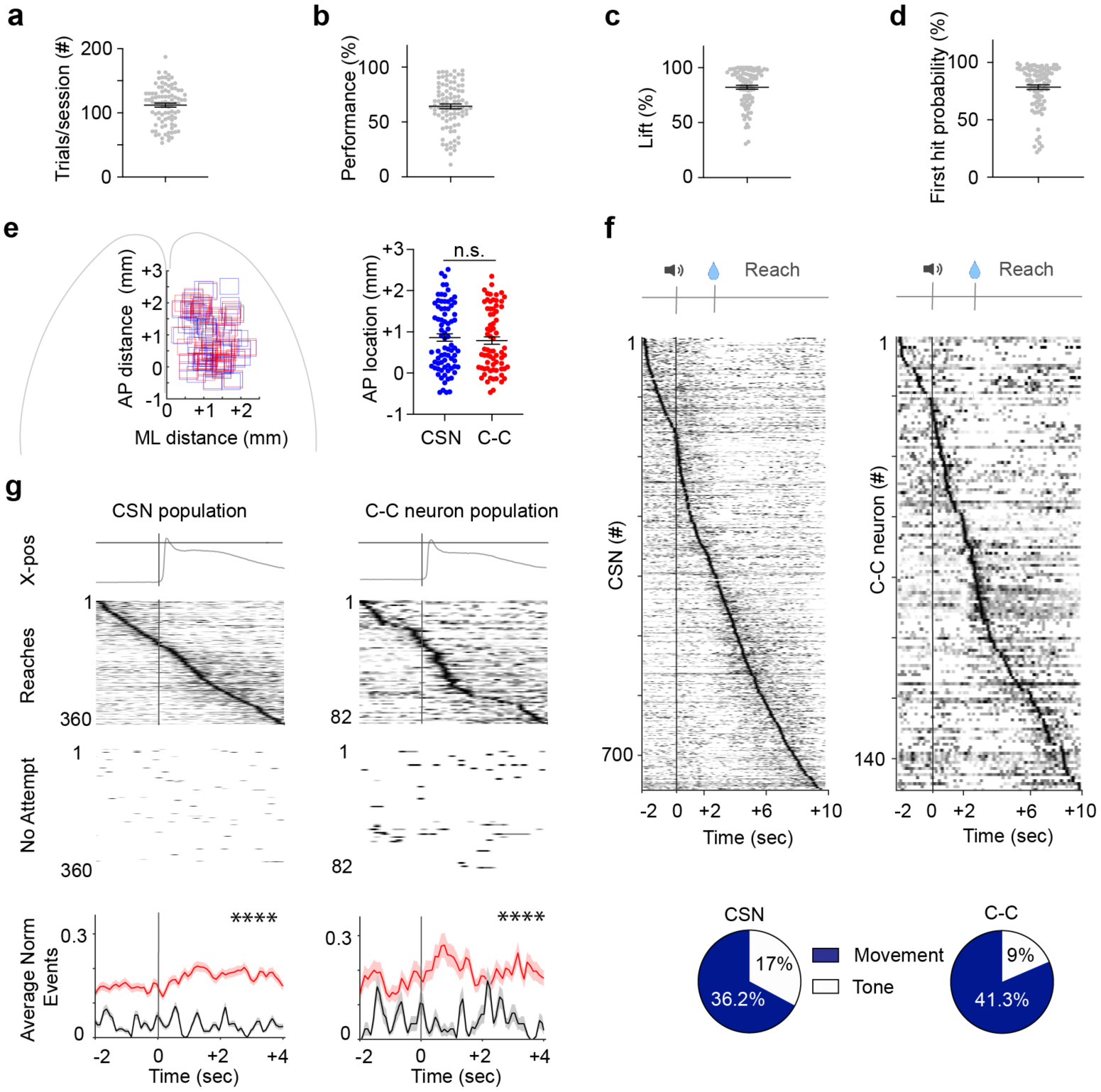
Quantification of reaching behaviour and 2-photon Ca^2+^ imaging. a. Well trained mice perform on average 112.1 ± 3.1 trials per session and day. b. The average performance rate (% hits per session) is 64. ± 2.2%. c. The average lift probability (% lifts per session) is 82.16 ± 1.79%. d. The first hit probability (% hits across all lifts in a session) is 78.49 ± 2.06%. e. The distribution of imaging FOVs for C-C neurons (red) and CSNs (blue) along the anterior-posterior (AP) and medio-lateral (ML) axis of the cortex. The FOVs for imaging C-C neurons and CSNs were distributed equally along the AP axis, Unpaired t-test, p = 0.5724. f. Task related modulation of C-C neurons and CSNs are aligned to tone, averaged and normalized. Cells are sorted by the time of peak modulation. Pie charts show the proportion of cells that modulate with tone or movement. g. The reach and no attempt trials for C-C neurons and CSNs. Both C-C neurons and CSNs had high firing rates during reaching and mostly silent in no attempt trials. Two-sample Kolmogorov-Smirnov test, p < 0.0001 for C-C neurons and CSNs.

